# N-Glycans on the extracellular domain of the Notch1 receptor control Jagged-1 induced Notch signalling and myogenic differentiation of S100β resident vascular stem cells

**DOI:** 10.1101/2023.11.17.567576

**Authors:** Eoin Corcoran, Abidemi Olayinka, Mariana di Luca, Yusof Gusti, Roya Hakimjavadi, Brendan O’Connor, Eileen M. Redmond, Paul A. Cahill

## Abstract

Notch signalling, critical for development and postnatal homeostasis of the vascular system, is highly regulated by several mechanisms including glycosylation. While the importance of O-linked glycosylation is widely accepted, the structure and function of N-glycans has yet to be defined. Here, we take advantage of lectin binding assays in combination with pharmacological, molecular, and site-directed mutagenetic approaches to study N-glycosylation of the Notch1 receptor. We find that several key oligosaccharides containing bisecting or core fucosylated structures decorate the receptor, control expression and receptor trafficking, and dictate Jagged-1 activation of Notch target genes and myogenic differentiation of multipotent S100β vascular stem cells. N-glycans at asparagine (N) 1241 and 1587 protect the receptor from accelerated degradation, while the oligosaccharide at N888 directly affects signal transduction. Conversely, N-linked glycans at N959, N1179, N1489 do not impact canonical signalling but inhibit differentiation. Our work highlights a novel functional role for N-glycans in controlling Notch1 signalling and differentiation of vascular stem cells.

**Teaser:** A sweet development in Notch regulation of vascular smooth muscle cell differentiation

## Introduction

The Notch signalling pathway is crucial for the development and postnatal homeostasis of the vascular system(*1*). It is activated by interaction of receptors (Notch1–4) and ligands (Delta-like 1, 3, 4, and Jagged 1, 2) present on the surface of adjacent cells (*2*). This interaction triggers sequential cleavage events resulting in release of the active form of Notch i.e., Notch intracellular domain (NICD). NICD translocates into the nucleus and, in combination with the transcription factor recombinant binding protein for the immunoglobulin region kJ (RBPJ, also known as CBF1/Suppressor of Hairless/LAG1, CSL) and the Mastermind-like (MAML 1–3) adaptor proteins, activates target genes including HES (Hairy and Enhancer of Split) and HEY (Hairy/enhancer-of-split related) that are essential modulators of neuronal, endocrine, and vascular cell fate (*3*). The Notch pathway may also be activated in the absence of ‘canonical’ ligands and independent of γ-secretase proteolytic cleavage; this non-canonical Notch signalling modulates Wnt/β-catenin, mammalian target of the rapamycin 2 complex (mTORC2)/Akt, Nuclear Factor kappa B (NF-κB), IkB kinase (IKK)-α/β and phosphatase and tensin homolog (PTEN)-induced kinase (PINK1)(*4*).

There is compelling evidence from lineage tracing studies for the involvement of (i) a rare subpopulation of differentiated medial vascular smooth muscle cells (SMC) that are stem cell antigen-1 (Sca1) positive (*5*, *6*) and (ii) various adventitial (*7*, *8*) and medial progenitors (*9–14*) in progressing subclinical atherosclerosis through the accumulation of ‘SMC-like’ cells within the vessel wall. Notch significantly impacts arterial pathology during lesion development by promoting SMC-like cell accumulation after injury (*12*, *15–18*). Multiple studies describe the ability of Notch signalling to induce myogenic differentiation of stem cell populations such as C3H 10T1/2 embryonic fibroblasts, mesenchymal stromal stem cells (MSCs) and neural crest stem cells (*19–21*). While the exact mechanism and cell(s) involved by which Notch and its target genes affect lesion formation remains incompletely understood, there is substantial evidence suggesting that the Notch1 receptor in particular plays a pivotal role in this process (*15–17*).

The Notch signalling pathway is tightly regulated by several mechanisms including cis-inhibition, negative feedback loops, PEST ubiquitination (*22*, *23*) and by glycosylation of the Notch receptor (*24*). With respect to the latter, the role of O-linked glycans has been well established (*25–27*). While there are also N-glycosylation sites present on the Notch receptor extracellular domain (ECD) (*28*), little is known about their specifics or their potential regulatory role in Notch signalling. The impact of N-glycan decoration of key proteins on a given process can be interrogated using chemical inhibitors of N-glycosylation. Tunicamycin (TNC), an antagonist for the GlcNAc phosphotransferase (GPT) enzyme, catalyses a critical step in the protein N-glycosylation pathway, linking the growing polypeptide to a dolichol precursor (*29*). 1-deoxymannojirimycin (DMJ) blocks the trimming of the 14-sugar dolichol precursor oligosaccharide by inhibiting the mannosidase I enzyme and preventing the synthesis of complex or hybrid-type N-glycans (*30*). Because the specificities of these inhibitors are broad, a more targeted approach for investigating N-glycans can be achieved using siRNA knockdown of key glycosyltransferases, Fut8 and Mgat3. A family of glycosyltransferases known as N-acetylglucosaminyl transferases (GnT’s) are responsible for the addition of GlcNAc residues to N-linked glycans. There are six members of this family (GnT-I to GnT-VI), all of which operate in the Golgi apparatus (*31*). GnT-III (coded by the *Mgat3* gene) catalyses the addition of a GlcNAc residue through a β-1,4 linkage to the β-mannose in the tri-mannosyl core of a growing N-linked oligosaccharide, creating a structure known as a bisecting N-glycan (*32*). This structure has been shown to inhibit the addition of sugar residues by other glycosyltransferases such as GnT-II, GnT-IV and Fut8 (*33*, *34*). Bisecting N-glycans regulate several processes, including cell adhesion, protein degradation and growth factor signalling (*35*). Members of the fucosyltransferase (FUT) family of enzymes facilitate the addition of fucose residues to the core N-glycan structure in a process known as core fucosylation (*36*). Fut8 catalyses the transfer of fucose from a donor substrate to the asparagine-149 bound GlcNAc at the base of an N-glycan via an α1,6-linkage (*37*). Core-fucosylated N-glycans regulate a wide array of cellular processes such as the expression of essential cadherins and the function of specific types of immunoglobulins (*38*, *39*). As of yet, it has not been established whether N-glycans on the Notch1 receptor are core fucosylated, though it has been noted that aberrant activity of both Fut8 and Notch is regularly observed in cancer (*40*).

Lectins have become a powerful experimental tool to determine the N-glycan profile of a glycoprotein, as they are highly specific carbohydrate-binding proteins with no catalytic activity (*41*). They can bind mono- or oligosaccharides and can recognise and distinguish between distinct branching and linkage patterns (*42*). Several lectin binding assays have been developed and optimised which exploit the high level of specificity seen in lectin-oligosaccharide binding to determine the glycosylation profile of a given glycoprotein. Unlike mass spectrometry and HPLC, these assays do not require the chemical or enzymatic removal of carbohydrates from proteins (*43*).

The aim of this study was to characterise the profile of N-glycosylation of the Notch1 receptor and explore the impact of these N-glycans on ligand induced Notch1 signalling, receptor trafficking and Notch-mediated myogenic differentiation of S100β vascular stem cells *in vitro*.

## Results

### Jagged-1 (Jag-1) activation of the Notch1 receptor promotes myogenic differentiation

Mouse aortic vascular stem cells (mVSCs) and C3H 10T1/2 embryonic fibroblasts were both positive for S100β and Sca1 expression, while negative for SMC markers Cnn1 and Myh11 [Fig S1A, Figure 1A]. S100β mVSCs exhibited increased telomere length and telomerase activity (indicators of stemness) when compared to aortic tissue and cultured SMCs [Fig S1B and C], and were capable of lineage specification following treatment with adipogenic and osteogenic differentiation media, respectively [Fig S1D and E].

**Figure 1:**
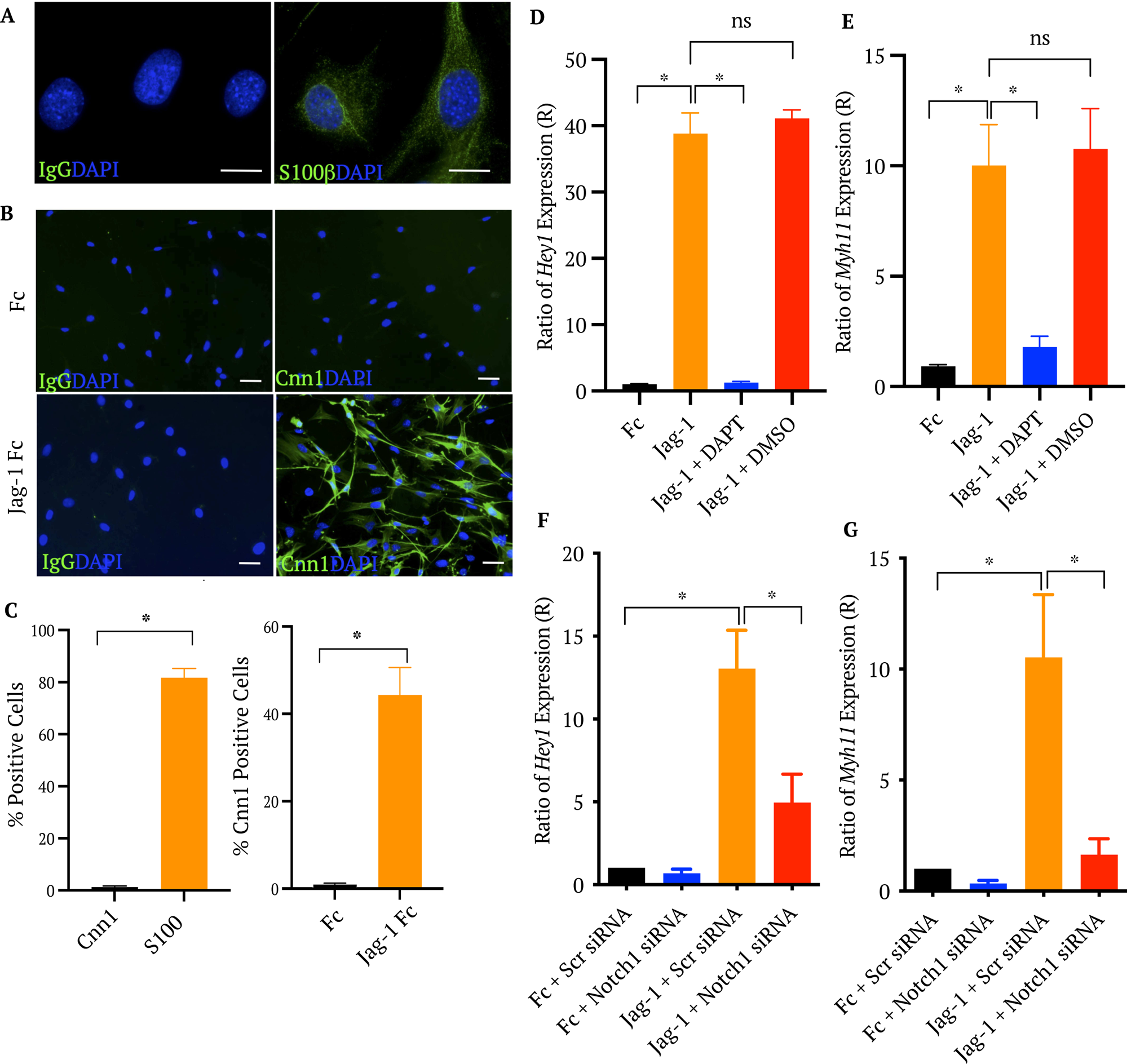
Jagged-1 activation of Notch1 signalling and SMC differentiation in S100β mVSCs. Immunocytochemical analysis of **A.** S100β protein expression in untreated aortic S100β mVSCs and **B.** Cnn1 expression in cells treated with recombinant immobilised Jagged-1 (Jag-1 Fc) or IgG-Fc at 1μg/mL in MM1 media after 7 days. Scale bar = 25μm. **C.** Mean percentage of S100β and Cnn1 positive cells. DAPI was used to stain nuclei. Positive cells were counted and compared against total DAPI cell count. Data are the mean ± SEM and representative of n=3 *p ≤ 0.05. **D-G.** Real-time qRT-PCR analysis of *Hey1* and *Myh11* mRNA levels in S100β mVSCs treated with recombinant immobilised Jagged-1 (Jag-1 Fc) or IgG-Fc (Fc) at 1μg/mL for 48 hrs with or without prior treatment of cells with a Y-secretase inhibitor, DAPT versus DMSO control (D, E) and following Notch1 receptor knockdown (F, G) versus scrambled control. The gene, hypoxanthine phosphoribosyltransferase *(hprt*) was used as a housekeeping control. Data are the mean ± SEM and representative of n=3 *p ≤ 0.05

Immobilised recombinant Jagged-1 (a Notch ligand) increased the number of SMC-like/myogenic Cnn1 positive cells (40.5 ± 5%, p<0.05 n=6) in S100β mVSCs cultures, when compared to the Fc-IgG control (0.1 ± 0.4%) [Figure 1B, C] concomitant with a significant increase in the expression of the Notch target gene, *Hey1* and the late SMC differentiation marker gene, *Myh11*, an effect attenuated following pre-treatment of the cells with the γ-secretase inhibitor, DAPT [Figure 1D and E].

Notch1 receptor depletion following siRNA knockdown at a transfection efficiency of > 80% [Fig S2A] reduced Notch1 protein expression [Fig S2B, C] and *Notch1* mRNA levels [Fig S2D] and significantly attenuated Jagged-1 induced *Hey1* [Figure 1F] and *Myh11* [Figure 1G] mRNA levels in S100β mVSCs, when compared to the scrambled control.

### Chemical inhibition of N-glycosylation impairs Jagged-1 induced Notch signalling

TNC and DMJ are both chemical inhibitors of N-glycosylation (*29*, *30*). N-glycan analysis was assessed in whole cell lysates and immunoprecipitated Notch1 samples from control, TNC and DMJ treated cells by measuring lectin binding using a series of commercial lectins (Table I). The binding characteristics of each lectin have previously been described (*43*).The optimal concentrations of TNC and DMJ required to reduce lectin binding in whole cell lysates was first evaluated [Figure 2A, B] before their effect on the N-glycan profile of the immunoprecipitated Notch1 receptor following ectopic expression was assessed [Figure 2C, D].

**Figure 2:**
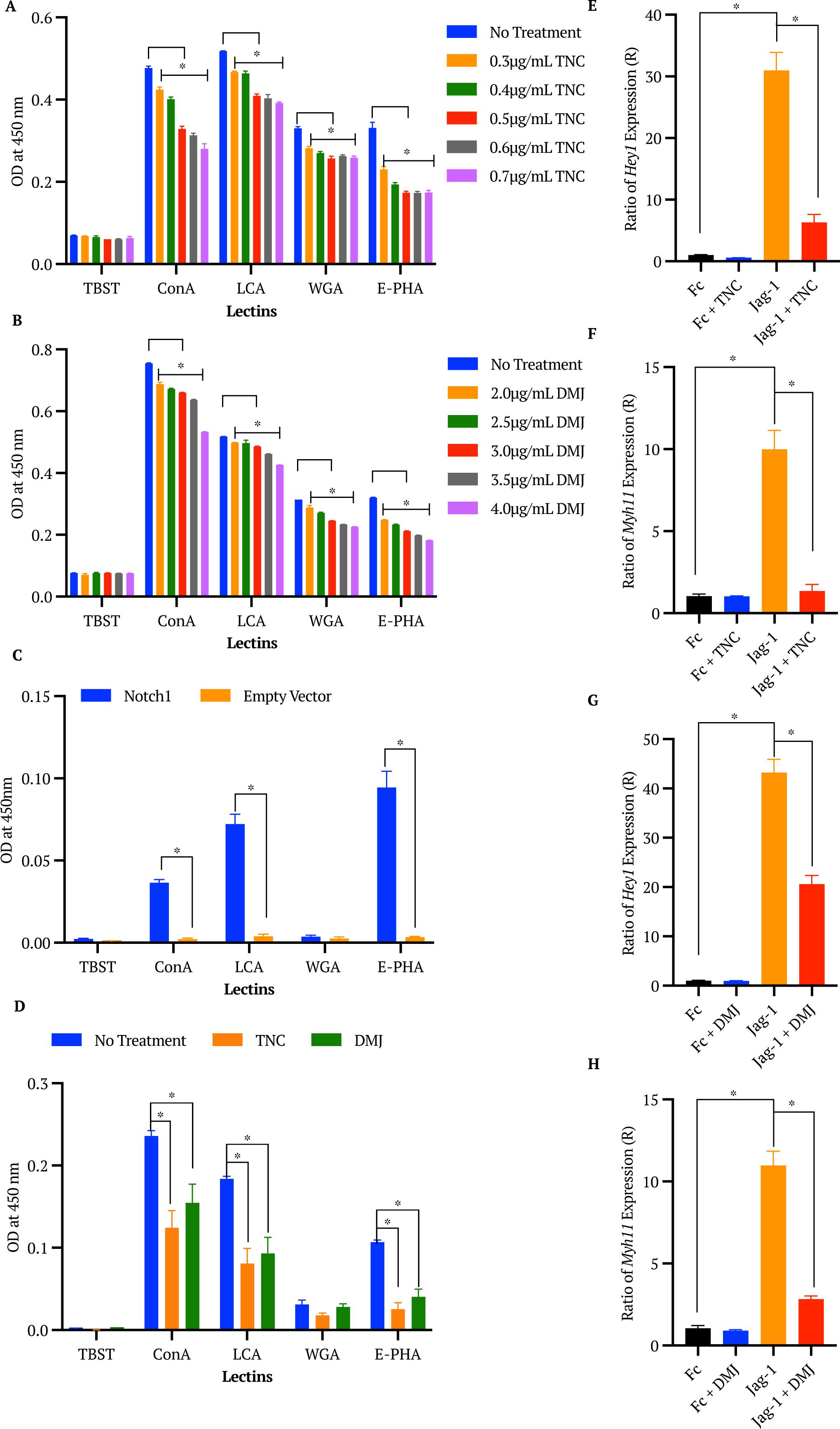
Chemical inhibition of N-glycosylation (TNC and DMJ) on lectin binding and Jagged-1 activation of Notch1 signalling and SMC differentiation in S100β mVSCs. **A-D.** Lectin binding analysis of N-glycans in whole cell lysates (A, C) and the immunoprecipitated Notch1 receptor following ectopic expression (B, D). S100β mVSC were treated with a range of TNC (A, B) and DMJ (C, D) concentrations for 48 h at 37°C. Data are presented as the mean absorbance value at 450 nm normalised to the negative control TBST. Data are the mean ± SEM and representative of n=3 *p ≤ 0.05 compared to the non-treated sample probed with the same lectins. **E-H.** Real-time qRT-PCR analysis of *Hey1* and *Myh11* mRNA levels in S100β mVSCs treated with recombinant immobilised Jagged-1 (Jag-1 Fc) or IgG-Fc (Fc) at 1μg/mL in MM1 media after 48 hrs following ectopic expression of the Notch1 receptor with or without prior treatment of cells with TNC (0.5μg/mL) (E, F) and DMJ (4μg/mL) (G, H). The gene, hypoxanthine phosphoribosyltransferase *(hprt*) was used as a housekeeping control. Data are the mean ± SEM and representative of n=3 *p ≤ 0.05

**Table I:**
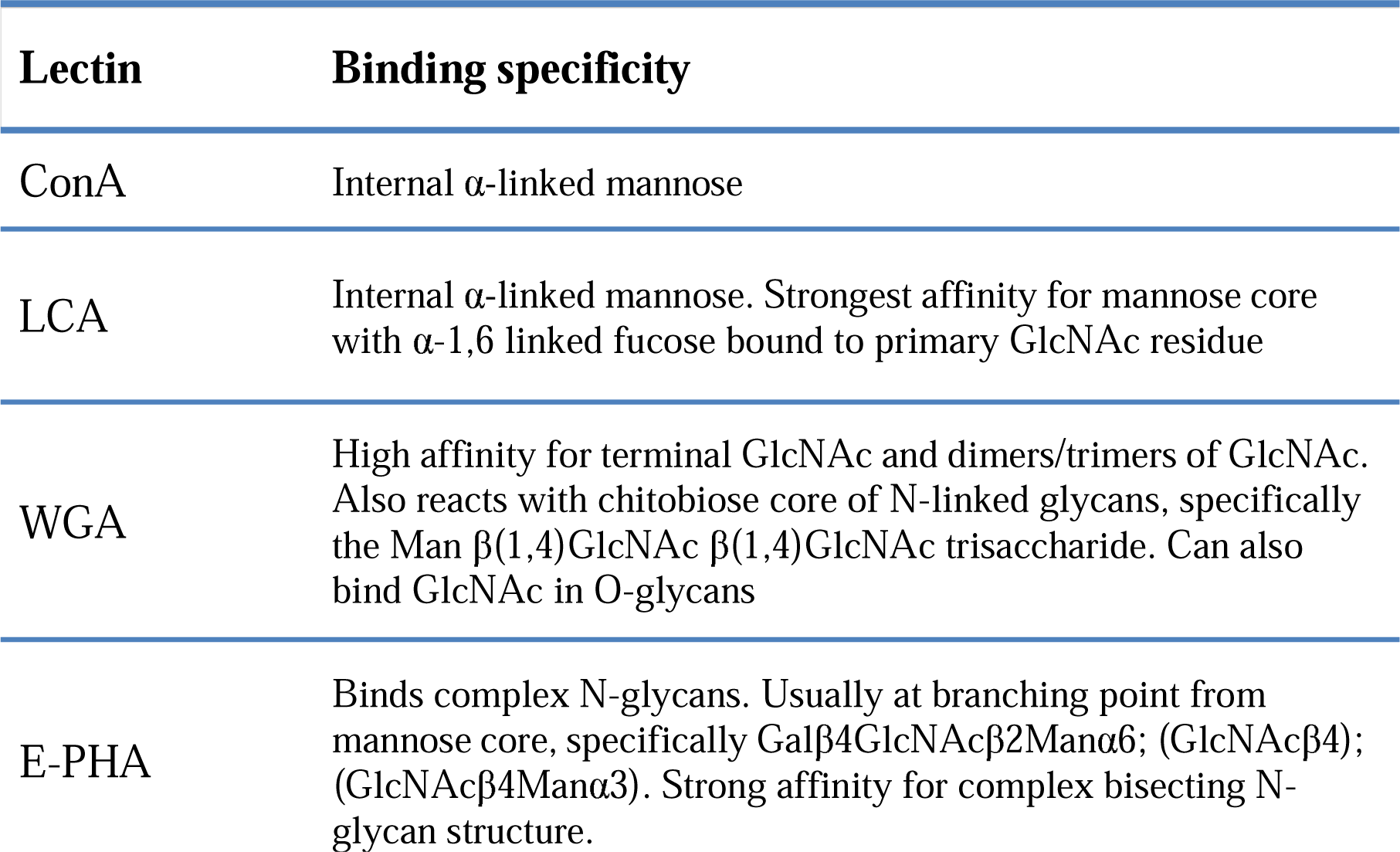
The binding specificities of commercial lectins. The binding specificity of Concanavalin A (ConA), Lens Culinaris Agglutinin (LCA), Wheat germ agglutinin (WGA) and Phaseolus vulgaris lectin E (E-PHA) on the ELLA assay.

| Lectin | Binding specificity |
| --- | --- |
| ConA | Internal α-linked mannose |
| LCA | Internal α-linked mannose. Strongest affinity for mannose core with α-1,6 linked fucose bound to primary GlcNAc residue |
| WGA | High affinity for terminal GlcNAc and dimers/trimers of GlcNAc. Also reacts with chitobiose core of N-linked glycans, specifically the Man $\beta$(1,4)GlcNAc $\beta$(1,4)GlcNAc trisaccharide. Can also bind GlcNAc in O-glycans |
| E-PHA | Binds complex N-glycans. Usually at branching point from mannose core, specifically Galβ4GlcNAcβ2Manα6; (GlcNAcβ4); (GlcNAcβ4Manα3). Strong affinity for complex bisecting N-glycan structure. |

There was a dose-dependent decrease in absorbance from whole cell lysates, indicating a reduction in lectin binding, following treatment of S100β mVSCs with TNC when compared to the non-treated control cells [Figure 2A]. Maximum inhibition was observed at 0.5μg/mL (in the cases of WGA and E-PHA). Similarly, DMJ treatment caused a significant reduction in binding across all lectins compared to the non-treated control with maximal inhibition observed at 4.0μg/mL [Figure 2B]. Lectin binding was significantly reduced in the immunoprecipitated Notch1 preparation from S100β mVSCs [Figure 2C] treated with both TNC [Figure 2D] and DMJ [Figure 2D] across all lectins, other than WGA. Inhibition by DMJ was as pronounced as that of TNC.

The Jagged-1 induced increase in *Hey1* and *Myh11* mRNA levels was significantly reduced in TNC [Figure 2E, F] and DMJ [Figure 2G, H] treated S100β mVSCs, when compared to untreated control cells. In parallel studies, ectopic expression of Notch1 NICD [Fig S3A, B] increased both *Hey1* and *Myh11* mRNA levels in S100β mVSCs [Figure 3A-D]. TNC and DMJ treatment inhibited the *Myh11* response [Figure 3B, D], but not the *Hey1* response [Figure 3A, C].

**Figure 3:**
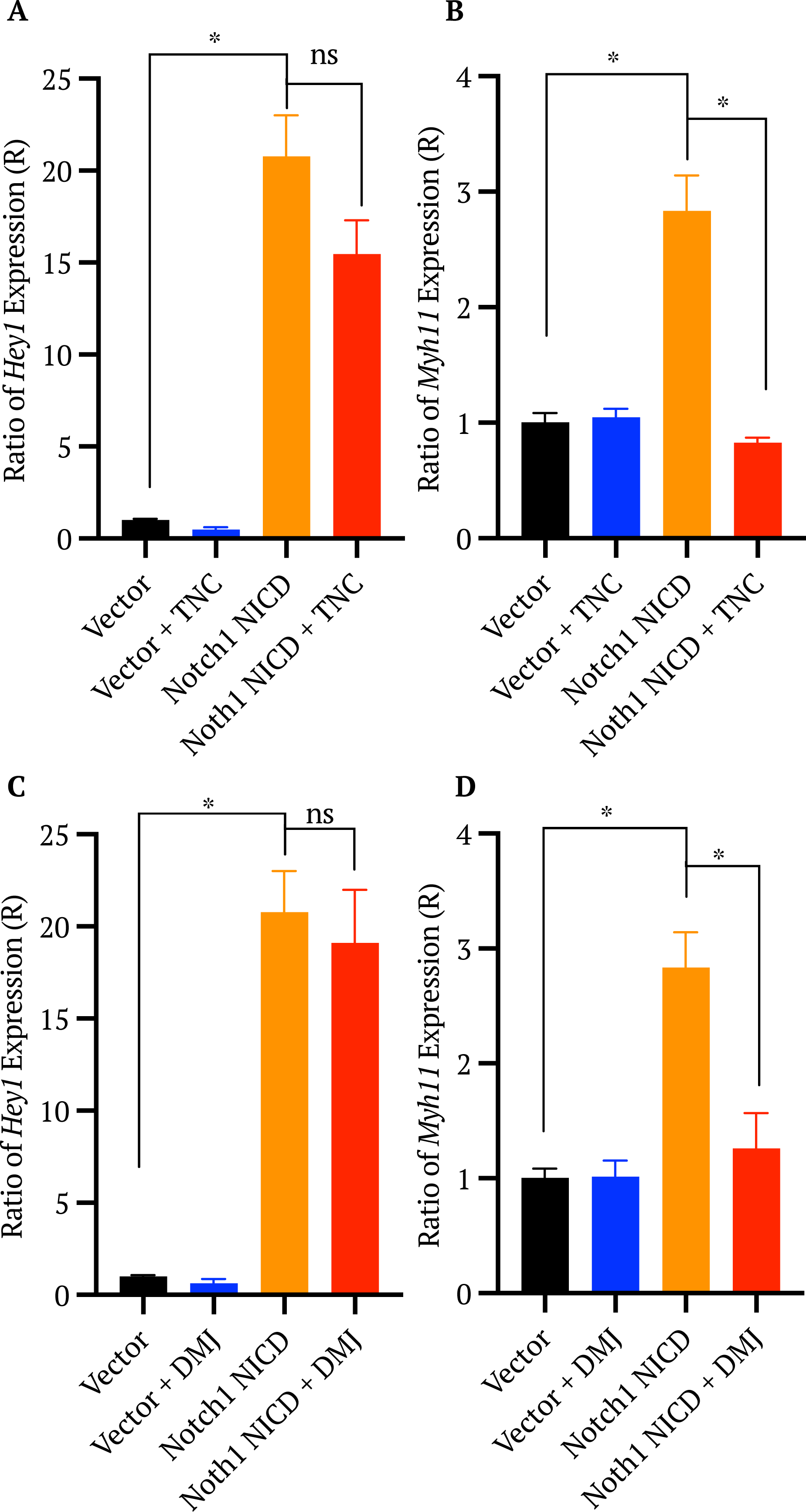
Chemical inhibition of N-glycosylation (TNC and DMJ) on Notch1 intracellular domain (NICD) activation of Notch signalling and SMC differentiation in S100β mVSCs. **A-D.** Real-time qRT-PCR analysis of *Hey1* (A, C) and *Myh11* (B, D) mRNA levels in S100β mVSCs following ectopic expression of the Notch1 NICD after 48 hrs with or without prior treatment of cells with TNC (0.5μg/mL) (A, B) or DMJ (4μg/mL) (C, D). The gene, hypoxanthine phosphoribosyltransferase *(hprt)* was used as a housekeeping control. Data are the mean ± SEM and representative of n=3 *p ≤ 0.05.

The reduction in Jagged-1 stimulated *Hey1* expression following TNC and DMJ treatment was not due to changes in the expression of the Notch1 receptor as there was no significant change in either *Notch1* mRNA levels or Notch1 protein expression following TNC or DMJ treatment [Fig S3C-E], compared to the non-treated control.

A similar dose-dependent decrease in lectin binding was observed in whole lysates following treatment of C3H10T1/2 cells with TNC [Fig S4A] and DMJ [Fig S4B], respectively but only for WGA and E-PHA lectins. No significant change in binding was detected at any DMJ concentration with ConA or LCA [Fig 4 SB]. This decrease in lectin binding in C3H10T1/2 cells following TNC and DMJ treatment was associated with a significant reduction in Jagged-1 induced in *Hey1* [Fig 4SD, F] and *Myh11* mRNA [Fig 4SE, G] levels, when compared to untreated control cells. Interestingly, TNC treatment also significantly decreased *Myh11* expression in the Fc-treated control sample [Fig S4 E].

These data suggest that global inhibition of glycosyltransferase activity reduces lectin binding of the Notch1 receptor, an effect concomitant with reduced ligand-induced Notch target gene expression and myogenic differentiation of S100β stem cells. This effect appears to be primarily at the level of the Notch1 receptor, as NICD-induced target gene expression was unaffected. However, NICD-induced myogenic differentiation is susceptible to inhibition following global removal of N-glycans.

### GnT-III and Fut8 depletion impairs Jagged-1 induced Notch signalling

*Mgat3* and *Fut8* genes code for the glycosyltransferases GnT-III and Fut8, respectively (*37*, *45*). To deplete their levels, targeted siRNA knockdown was performed in S100β mVSCs [Fig S3F, G] and C3H10T1/2 cells [Fig S5A, B] before the N-glycan profile of whole lysates [Figure 4A] and immunoprecipitated Notch1 [Figure 4B] was assessed for lectin binding using the ELLA assay. Lectin binding to ConA and E-PHA was reduced in whole cell lysates following *Mgat3* knockdown in both S100β mVSCs [Figure 4A] and C3H10T1/2 cells [Fig S4C] compared to the scrambled control, with no effect on LCA or WGA lectins. *Fut8* knockdown led to a significant decrease in LCA binding only [Figure 4A, Fig S4C]. Similarly, lectin binding of immunoprecipitated Notch1 following *Mgat3* knockdown was reduced in S100β mVSCs for both ConA and E-PHA lectins, compared to the scrambled control, while no reduction in binding was detected with LCA or WGA lectins [Figure 4B]. *Fut8* knockdown led to significant decreases in ConA and LCA binding, but not for WGA or E-PHA lectins [Figure 4B].

**Figure 4:**
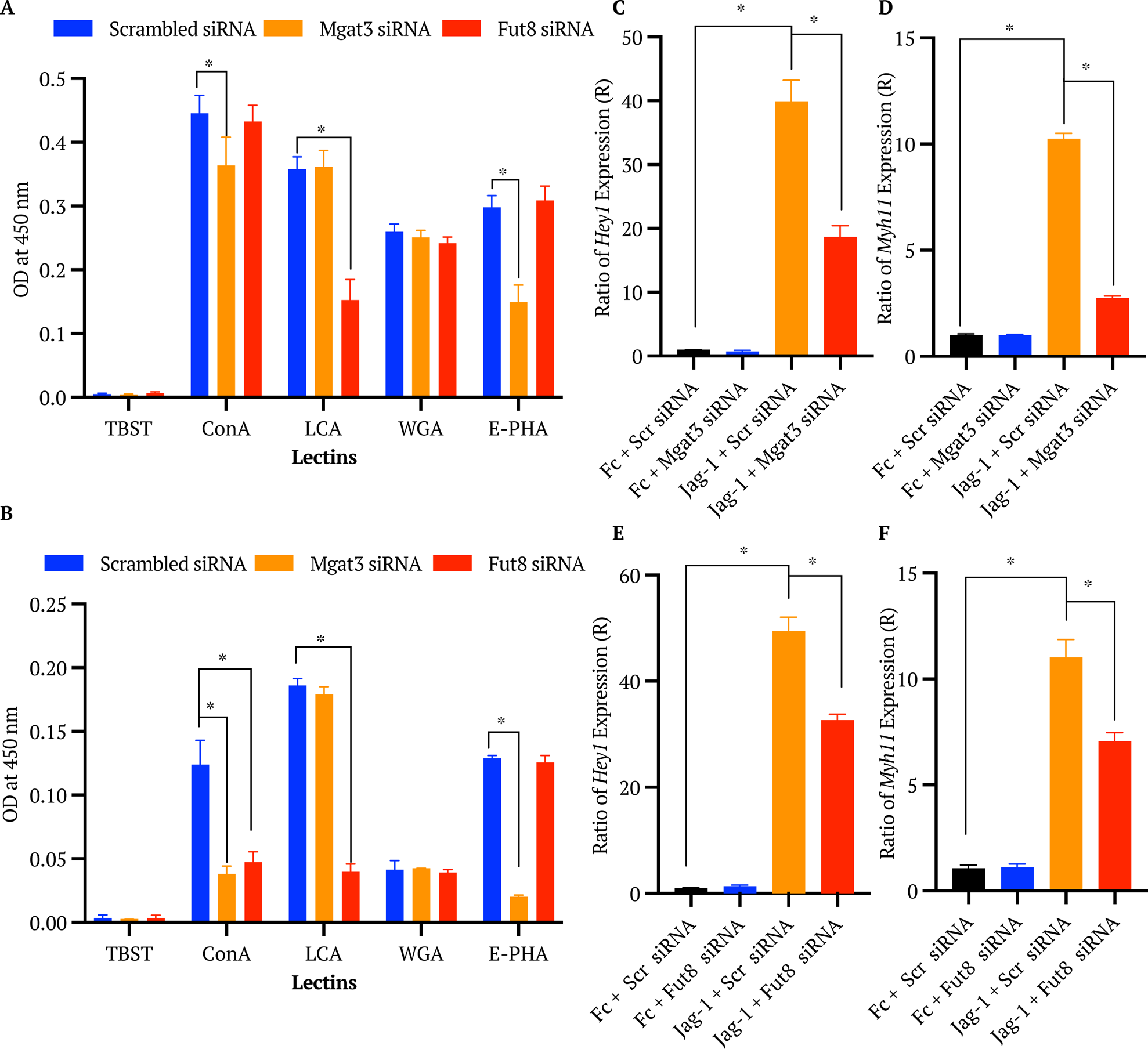
*Mgat3* and *Fut8* knockdown on lectin binding, Notch signalling, and SMC differentiation in S100β mVSCs. **A, B.** Lectin binding analysis of N-glycans in whole cell lysate (A) and the immunoprecipitated ectopic Notch1 receptor (B) following *Mgat3* and *Fut8* knockdown with specific siRNA duplexes (25nM) versus scrambled control. Data are presented as the mean absorbance value at 450 nm normalised to the negative control TBST sample. Data are the mean ± SEM and representative of n=3 *p ≤ 0.05. compared to the non-treated sample probed with the same lectins. **C-F.** Real-time qRT-PCR analysis of *Hey1* (C, E) and *Myh11* (D, F) mRNA levels in S100β mVSCs treated with recombinant immobilised Jagged-1 (Jag-1 Fc) or IgG-Fc (Fc) at 1μg/mL after 48 hrs following *Mgat3* (C, D) or *Fut8* (E, F) knockdown with specific siRNA duplexes (25nM) versus scrambled control. The gene, hypoxanthine phosphoribosyltransferase *(hprt*) was used as a housekeeping control. Data are the mean ± SEM and representative of n=3 *p ≤ 0.05.

Jagged-1 stimulated *Hey1* and *Myh11* expression was reduced following *Mgat3* and *Fut8* knockdown in both S100β mVSCs [Figure 4C-F] and C3H10T1/2 cells [Fig S5C-F], when compared to the scrambled controls. In contrast, Notch1 NICD-induced *Hey1* and *Myh11* mRNA levels were unaffected by *Mgat3* [Figure 5A, B] and *Fut8* knockdown [Figure 5C, D] in S100β mVSCs. Moreover, the decrease in *Hey1* expression following *Fut8* knockdown was not due to changes in the expression of the Notch1 receptor as there was no significant change in either Notch1 protein expression [Figure 5E, F] or *Notch1* mRNA levels [Figure 5G] following knockdown, compared to scrambled control. In contrast, while there was a significant change (albeit slight) in the expression of Notch 1 protein levels following *Mgat3* knockdown [Figure 5E, F] this was not reflected in changes to *Notch1* mRNA levels [Figure 5G].

**Figure 5:**
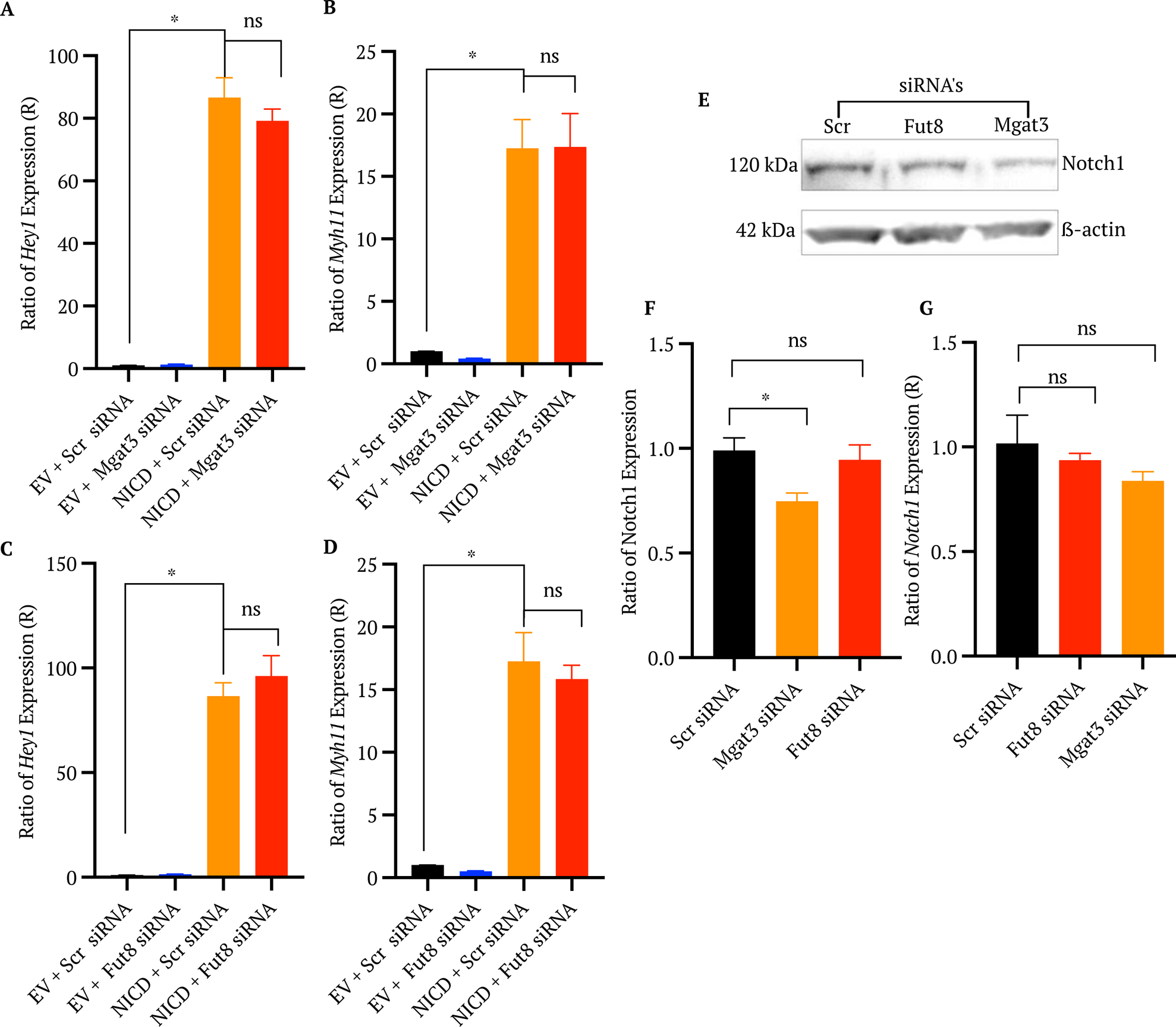
*Mgat3* and *Fut8* knockdown on Notch1 intracellular domain (NICD) activation of Notch signalling and SMC differentiation in S100β mVSCs. **A-D.** Real-time qRT-PCR analysis of *Hey1* (A, C) and *Myh11* (B, D) mRNA levels in S100β mVSCs following ectopic expression of Notch1 NICD after *Mgat3* (A, B) and *Fut8* (C, D) knockdown with specific siRNA duplexes (25nM) compared to the scrambled (*Scr*) siRNA control. **E-F.** Notch1 protein levels and *Notch1* mRNA levels in whole cell lysates from S100β mVSCs following *Mgat3* or *Fut8* knockdown with specific siRNA duplexes (25nM) versus scrambled control. **E.** A representative immunoblot image of Notch1 bands at 120kDa and β-actin bands at 42kDa. **F.** densitometry analysis of the intensity of Notch1 bands normalised to the β-actin loading control. Notch1 protein levels presented as the ratio of expression relative to the untreated negative control sample. **G.** Real-time qRT-PCR analysis of *Notch1* mRNA levels following *Mgat3* or *Fut8* knockdown with specific siRNA duplexes (25nM) versus scrambled control. The gene, hypoxanthine phosphoribosyltransferase *(hprt*) was used as a housekeeping control. Data are the mean ± SEM and representative of n=3, ns or *p ≤ 0.05.

### Inhibition of N-glycosylation impairs Notch receptor trafficking and degradation

N-glycosylation is an essential modification for correct protein trafficking from the endoplasmic reticulum (ER) and Golgi apparatus (GA) to their site of action within the cell (*24*, *44*). Therefore, the possibility that inhibition of N-glycosylation interferes with Notch1 protein trafficking and localisation to the membrane was explored. Membrane fractionation of crude lysate from S100β mVSCs revealed the presence of the Notch1 receptor in the membrane fraction, confirmed using pan-cadherin, a ubiquitously expressed membrane-bound protein [Figure 6A]. Following treatment of the cells with TNC or DMJ for 48 h, there was a significant reduction in the expression of Notch1 in the membrane fraction [Figure 6B, D]. Brefeldin A (BFA), as a potent inhibitor of protein transport in the Golgi apparatus, served as a positive control. Similarly, there was a significant reduction in the expression of Notch1 within the membrane fraction following *Mgat3* knockdown, compared to the scrambled control [Figure 6C, E]. In contrast, there was no significant change following *Fut8* knockdown [Figure 6C, E]. This analysis suggests that Notch1 trafficking from the GA to the plasma membrane is inhibited by removal of a bisecting N-glycan.

**Figure 6:**
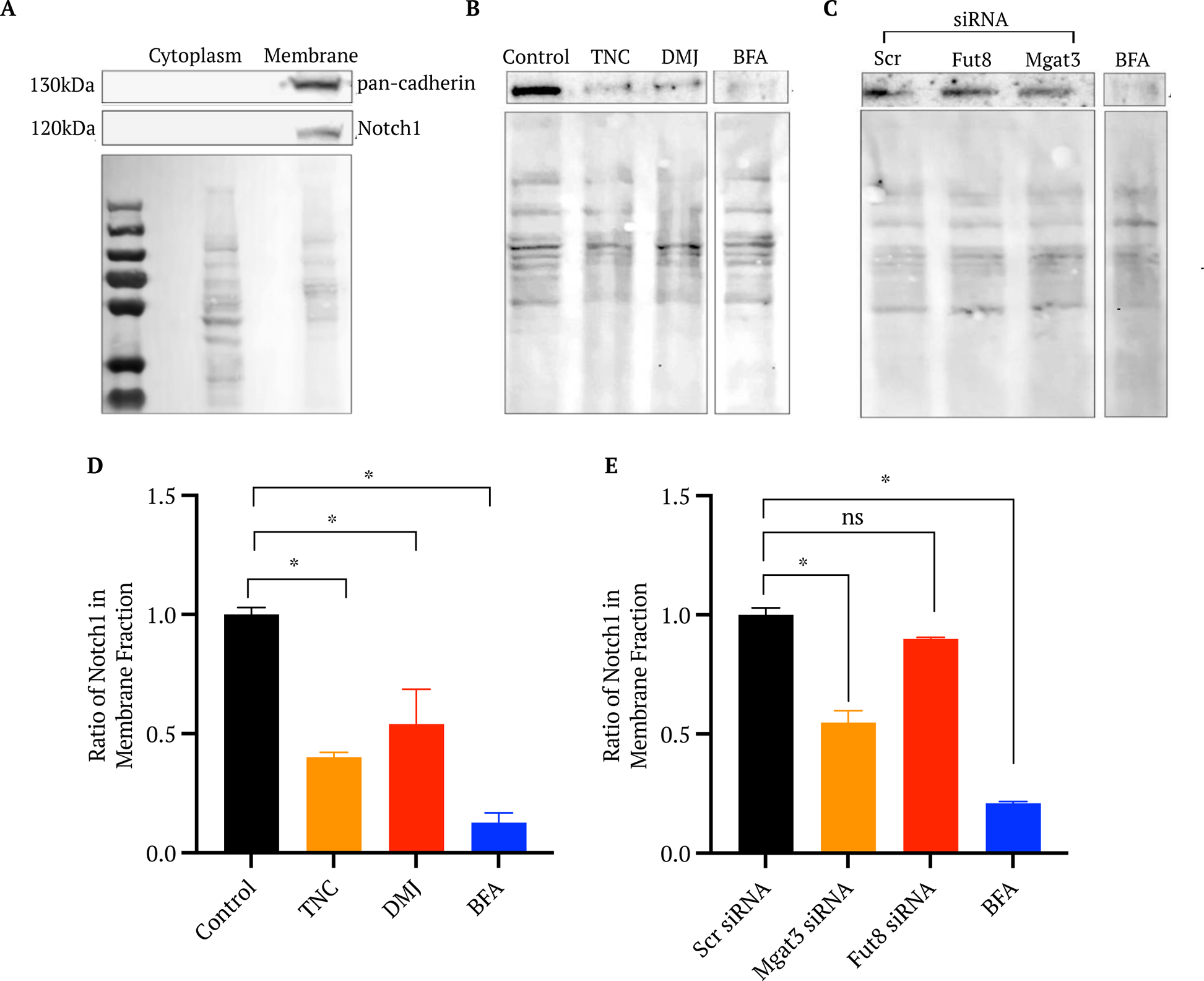
Chemical inhibition of N-glycosylation (TNC and DMJ) and *Mgat3* and *Fut8* knockdown on membrane localisation of the Notch1 receptor. **A.** Notch1 and Pan-Cadherin protein expression in cytoplasmic and membrane fractions from S100β mVSCs lysates. A representative immunoblot image of Notch1 at 120kDa (upper panel) and Ponceau S staining (lower panel) in each fraction. **B.** Notch 1 protein expression in the membrane fraction of whole cell lysates from S100β mVSCs treated with TNC (0.5μg/mL), DMJ (4μg/mL) or 1 X BFA for 48 h at 37°C. **C.** Notch 1 protein expression in the membrane fraction of whole cell lysates from S100β mVSCs following *Mgat3* or *Fut8* knockdown compared to scrambled control or cells treated with 1 X BFA. **D, E.** Densitometry analysis of the intensity of Notch1 bands normalised to the Ponceau S loading control. Notch1 protein levels in membrane fraction presented as the ratio of expression relative to the untreated negative control or scrambled control. Data are the mean ± SEM and representative of n=3, ns or *p ≤ 0.05.

These data were further corroborated by analysing Notch1 receptor localisation in cells before and after inhibition of N-glycosylation and determining the ratio of membrane:perinuclear staining *in situ* using immunocytochemistry. Lower ratios indicate a higher level of fluorescence in the perinuclear region compared to that at the membrane. Golgin97 served as a control to visualise the subcellular location of the GA. The ratio of both the fluorescence density and intensity significantly decreased following treatment with TNC, DMJ or BFA wherein a significant decrease in the ratio of membrane:perinuclear localisation of Notch1 was observed following treatment [Figure 7A, B, C]. Collectively, these data suggest an accumulation of Notch1 in the perinuclear region of the cell in response to TNC and DMJ treatment, similar to BFA and supports reduced Notch1 protein trafficking following inhibition of N-glycosylation. Since cellular Notch1 protein levels remained unchanged following treatment with TNC and DMJ [Fig S3D, E], it is unlikely that TNC and DMJ treatment promotes endocytosis and degradation of the receptor.

**Figure 7:**
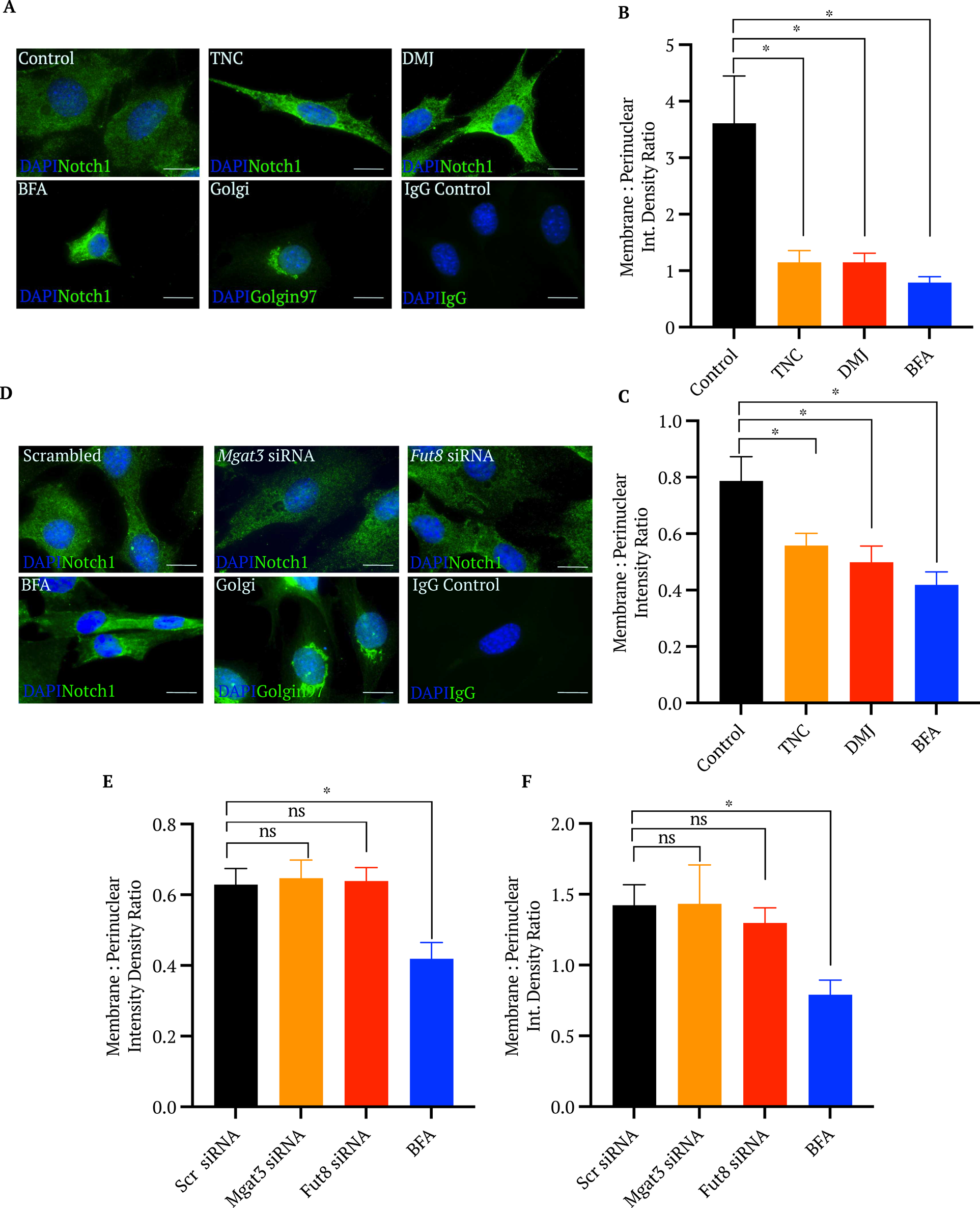
Chemical inhibition of N-glycosylation (TNC and DMJ) and *Mgat3* and *Fut8* knockdown on Notch1 receptor trafficking. **A.** Immunocytochemical analysis of the localisation of the Notch 1 receptor in S100β mVSCs treated with TNC (0.5μg/mL), DMJ (4μg/mL) or 1 X BFA. **B, C.** ImageJ analysis of the ratio of Membrane:Perinuclear fluorescence using (B) integrated density and (C) mean fluorescence intensity values. Representative image at 60X magnification. Location of the GA was confirmed using Golgin97. Scale bar = 25μm. Data are the mean ± SEM and representative of n=3, ns or *p ≤ 0.05. **D.** Immunocytochemical analysis of the localisation of the Notch 1 receptor in S100β mVSCs following *Mgat3* or *Fut8* knockdown. (D) Representative image at 60X magnification. **E, F.** ImageJ analysis of the ratio of Membrane:Perinuclear fluorescence using (B) integrated density and (C) mean fluorescence intensity values Location of the GA was confirmed using Golgin97. Scale bar = 25μm. Data are the mean ± SEM and representative of n=3, ns or *p ≤ 0.05.

In contrast, there was no significant change in the ratio of either the fluorescence density or intensity following *Mgat3* or *Fut8* knockdown whereas both ratios decreased significantly following BFA treatment [Figure 7D, E, F]. Taken together, these data suggest that despite a reduction in the expression of Notch1 in the membrane fraction following *Mgat3* knockdown, it is more likely that this reduction is due to the decrease in total Notch1 protein in the cell following *Mgat3* knockdown [Figure 5E, F].

### Site-directed mutagenesis of six N-glycosylation impairs Jagged-1 induced Notch signalling

To further define the function of both bisecting and core fucosylated N-glycans, site-directed mutagenesis of the six N-glycosylation consensus sequences along the Notch1 ECD was performed. Site-directed mutagenesis of plasmid-encoded Notch1 was chosen, and the proportion of transfected cells was maximised in each population through puromycin selection [Fig S6A, B]. The optimum concentration of plasmid DNA was assessed using a green fluorescent protein (GFP)-coding plasmid [Fig S6B]. Using the *pCS2 Notch1 Full Length-6MT* plasmid, six single base substitutions were carried out separately on the Notch1 gene to alter asparagine residues at amino acid positions 888, 959, 1179, 1241, 1489 and 1587 to isoleucine residues. Mutant plasmids were sequenced and aligned with the wild-type plasmid sequence using Multalin to confirm that the correct single base substitution had been incorporated in each case. In each case an essential adenine base was correctly converted to a thymine base [Fig S7A]. Chromatograms were also generated to determine the quality of both the sequencing read and the sample plasmid and confirmed high quality with clearly defined peaks [Fig S7B].

Mutations at N888I, N1489I and N1587I all resulted in decreased ConA, LCA and E-PHA binding, compared to that observed in the wild-type cells, with no effect on WGA binding. N959I led to a significant reduction only in ConA binding, while N1179I caused a significant attenuation in LCA binding only [Figure 8A]. In parallel cultures, Jagged-1 significantly increased the expression of *Hey1* in control wild-type Notch1 transfected cells, an effect that was significantly attenuated by mutations at N888I, N1241I and N1587I [Figure 8B] concomitant with a significant reduction in Jagged-1 induced *Myh11* expression across all mutants [Figure 8C].

**Figure 8:**
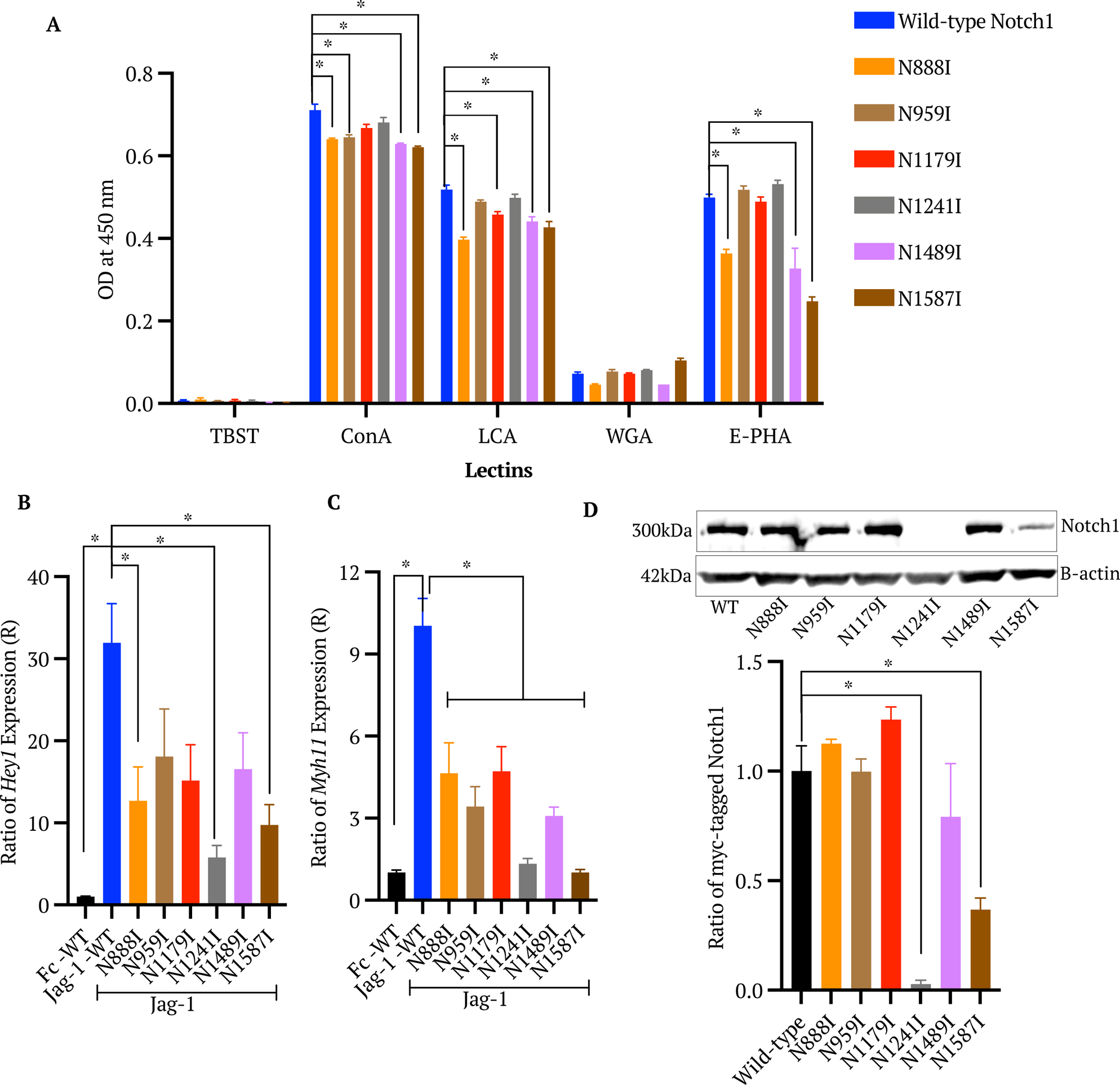
Site-directed mutagenesis of asparagine residues on the Notch1 ECD receptor on lectin binding, Notch signalling, and SMC differentiation in C3H10T1/2 cells. **A.** Lectin bindings analysis of N-glycans on immunoprecipitated wild-type and mutant Notch1 receptors (N888I, N959I, N1179I, N1241I, N1489I and N1587I) from puromycin selected C3H10T1/2 cells. Data are presented as the mean absorbance value at 450nm normalised to the negative control TBST sample and the IgG immunoprecipitation control. **B, C.** Real-time qRT-PCR analysis of Jag-1 stimulated *Hey1* (B) and *Myh11* (C) mRNA levels in C3H10T1/2 cells following ectopic expression of wild-type and mutant Notch1 receptors. Cells were treated with recombinant immobilised Jagged-1 (Jag-1 Fc) or IgG-Fc (Fc) at 1μg/mL in MM1 media after 48 hrs. The gene, hypoxanthine phosphoribosyltransferase *(hprt*) was used as a housekeeping control. **D.** Immunoprecipitated wildtype (WT) and mutant Notch1 receptor expression. A representative immunoblot image of full-length Notch1 bands at 300kDa and β-actin bands at 42kDa. Densitometry analysis of the intensity of Notch1 bands was normalised to the β-actin loading control. Full-length Notch1 protein levels presented as the ratio of expression relative to the wild-type Notch1 control. Data are the mean ± SEM and representative of n=3, ns or *p ≤ 0.05.

The changes in *Hey1* or *Myh1*1 expression were not attributed to differences in plasmid transfection efficiency across mutants as there was no significant difference between levels of Notch1-coding plasmid DNA across all mutant plasmid samples compared to the wild-type control [Fig S8A]. However, a decrease in Notch1 protein levels was observed in cells transfected with mutants N1241I and N1587I, respectively, while no change was detected across all other mutants, compared to the wild-type control [Figure 8D]

These data indicate that all the mutations incorporated into the Notch1 gene, except N1241, had a measurable effect on receptor glycosylation. Lectin binding indicated the presence of a range of different N-glycan structures on the Notch1 ECD, including core fucosylated and bisecting N-glycans, and some additional structures that could not be identified with the limited range of lectins used. Mutation of asparagine residues at Notch1 N-glycosylation sites reduced *Hey1* expression (N888I, N1241I and N1587I) and *Myh11* expression (all mutations) following Jagged-1 stimulation. This was not due to altered plasmid transfection efficiency, but for some mutants (N1241I and N1587I), altered expression and/or degradation of Notch1 may be involved. It is likely that the N-glycans at the four other mutation sites may influence Notch1 processing and signalling, such as receptor cleavage or trafficking.

## Discussion

In contrast to the well documented role of O-glycosylation, very little is known about the structure and regulatory function of N-linked glycans and their impact on Notch signalling. Here we report on bisecting and core fucosylated N-glycans decorating the extracellular domain (ECD) of the Notch1 receptor and demonstrate their role in regulating ligand-induced Notch signalling and myogenic differentiation of vascular stem cells.

Notch drives myogenic differentiation of a range of stem cells, including S100β resident mVSCs(*19*, *20*, *45*). Herein, immobilised recombinant Jagged-1 was used to activate Notch signalling and stimulate the expression of both Notch target genes and myogenic genes in S100β mVSCs. To date, evidence regarding the profile of N-linked glycans on the Notch1 receptor suggests that six consensus sites are present and highly conserved along the ECD, and that at least some of the N-glycans at these sites contain a bisecting structure (*28*, *46*). In the current study, lectin binding analysis of the purified Notch1 receptor confirmed the presence of N-glycans, and of bisecting N-glycans, whilst also revealing the existence of oligosaccharides with a fucose residue bound to the core GlcNAc by an α-1,6 linkage (referred to as core fucosylation). Global N-glycosylation inhibition with tunicamycin (TNC) and 1-deoxymannojirimycin (DMJ) further supported these findings as each inhibitor promoted a reduction in binding of E-PHA (specific for bisecting structures) and LCA (specific for core fucosylated structures) lectins to the purified Notch1 protein. However, as many lectins have weak affinity for oligosaccharide structures other than their main binding target, including both E-PHA and LCA (*47*, *48*), knockdown of the key glycosyltransferases responsible for the formation of bisecting and core fucosylated N-glycans, namely GnT-III and Fut8, confirmed their presence on the Notch1 ECD.

Knockdown of the *Mgat3* and *Fut8* glycosyltransferase-coding genes had a significant effect on bisecting or core fucosylated N-glycans of Notch1, supporting the TNC and DMJ inhibition studies. *Mgat3* and *Fut8* knockdown also reduced ligand-mediated Notch signalling and myogenic differentiation in both mVSCs and C3H 10T1/2 cells. As downstream components of Notch signalling and myogenic differentiation pathways following ectopic expression of Notch1 NICD were not altered by *Mgat3* or *Fut8* knockdown, their effect appears to be primarily at the level of the Notch1 receptor. Furthermore, as Notch1 transcript levels were not affected by either knockdown, but Notch1 protein expression was significantly reduced by *Mgat3* expression, these data suggest that bisecting N-glycans may have a role in preventing the degradation of the Notch1 receptor. The function of core fucosylated N-glycans is more difficult to decipher as reduction of their synthesis through *Fut8* knockdown had no effect on Notch1 protein expression or on receptor trafficking but caused a significant attenuation of Notch signalling and myogenic differentiation. There may be an alternative function for core fucosylated N-glycans on the Notch1 receptor, such as regulating cleavage events. Our finding that inhibition of GnT-III blocks Notch activation suggests that targeting GnT-III or blocking bisecting glycosylation sites on the Notch1 receptor may lead to new therapeutics targeting neointimal formation. Overall, the evaluation of GnT-III expression levels could also have clinical utility for stratifying patient cohorts that could benefit from therapies that target Notch1 signalling.

While the importance of O-linked glycosylation in Notch receptor-ligand binding and receptor processing and trafficking is well documented (*25*, *27*, *49*), little information is available on the role of N-glycans in this context. It is clear that global inhibition of N-glycosylation following TNC and DMJ led to an accumulation of the receptor in the ER-Golgi complex concomitant with an attenuated Jagged-1 induced response in these cells suggesting that it was mediated in part by preventing Notch1 receptor trafficking out of the ER-Golgi complex to the plasma membrane. A similar effect of TNC on Notch 3 trafficking has also been reported (*50*). While this may be due to changes in glycosylation of the numerous proteins involved in receptor trafficking, several previous reports have demonstrated that inhibiting N-glycosylation can block their transport to the membrane (*51*, *52*). Therefore, it is likely that some of the N-glycans on the Notch1 ECD are required for correct trafficking of the protein to the plasma membrane. Unlike TNC and DMJ treatment, knockdown of *Mgat3* and *Fut8* did not affect downstream elements of either Notch signalling or myogenic differentiation pathways, indicating that bisecting and core fucosylated N-glycans exert their influence on the Jagged-1 response either at, or before, the point of Notch receptor activation by the ligand. Indeed, analysis of receptor trafficking suggests that inhibition of the glycosyltransferases did not result in accumulation of Notch1 in the ER-Golgi complex. This suggests that the inhibition of trafficking observed following TNC and DMJ treatment was either due to altered glycosylation of transport proteins or removal of N-glycans on the Notch1 receptor which do not contain bisecting or core fucosylated structures. A combination of the receptor trafficking data and measurement of Notch1 protein levels in the whole cell lysates and in the membrane compartment suggests that bisecting N-glycans may prevent the accelerated degradation of the Notch1 receptor. This supports previous studies showing Notch1 receptor colocalization with lysosomal degradation proteins following *Mgat3* knockdown (*53*). The potential impact of N-linked glycans on O-fucose glycosylation of Notch is unclear. N-linked sites within Notch1 ECD are in close proximity to O-fucose modified sites and O-Fucose glycosylation of Notch EGF repeats either inhibits or activates Notch ligand-dependent activation (*54*). It will be important in the future to determine how the bisecting N-glycan structures on the Notch1 receptor in mVSCs may alter this O-fucose code. The role of core fucosylated N-glycans may be more directly related to ligand activation of the Notch1 receptor as knockdown of the *Fut8* glycosyltransferase inhibited Notch signalling and myogenic differentiation but did not affect Notch1 expression, trafficking, degradation, or NICD downstream signalling.

Collectively, these data suggest a putative role for N-linked bisecting glycosylation in ligand activation of Notch1 in S100β mVSCs. However, as both chemical inhibition and knockdown of *Mgat3* and *Fut8* could also alter the N-glycosylation of proteins other than the Notch1 receptor, it is possible that these interventions could influence Notch downstream events. Indeed, nicastrin, for example, part of the γ-secretase complex which cleaves the NICD from the Notch1 transmembrane domain (TMD) following receptor activation, contains bisecting N-glycans which may be crucial for its role in the cleavage process (*46*, *55*). Similarly, *Fut9* knockdown of core fucosylated N-glycans impairs the proliferation of S100β neural stem cells via inhibition of Musashi-1 (Msi-1), an RNA-binding protein and activator of the Notch signalling pathway, that binds to *Numb* mRNA and inhibits its translation. *Fut9* knockdown is coupled with increased Numb levels eventually resulting in the suppression of Notch signalling (*56*). Therefore, the reduction in Notch target gene and myogenic gene expression following *Mgat3* and *Fut8* knockdown may have been due, at least in part, to removal of N-glycans (bisected and core fucosylated) from other key proteins impacting on Notch signalling in these cells. For this reason, it was important to further explore the mechanisms by which core fucosylated and/or bisecting N-glycans affected the Notch signalling process following site-directed mutagenesis of key sites on the Notch1 ECD.

Site-directed mutagenesis of the six N-glycosylation consensus sequences along the Notch1 ECD defined the function of both bisecting and core fucosylated N-glycans. While initial analysis indicated that the amino acid site N1241 may not be glycosylated, additional information suggested that it is more likely that the N-glycan at this site was not detected due to the limited range of lectins used and the low binding affinity of some lectins *in vitro*(*57*). This is further corroborated by the fact that the N1179 mutation did not lead to a significant change in ConA binding, but did significantly reduce the level of LCA binding, indicating that a core fucosylated oligosaccharide may be present at this site which was not detected by ConA. Similarly, analysis of the N959 mutant indicated that an N-linked oligosaccharide is present at this site but does not contain structures required for binding to LCA, WGA or E-PHA. Meanwhile, the glycan at site N1179 was found to contain a core fucosylated structure. Finally, it is noteworthy that N-glycans at N888, N1489 and N1587 shared a similar lectin binding profile with significant reduction in the level of binding with ConA, LCA and E-PHA with these mutants. Previous studies have demonstrated that GnT-III modification of an oligosaccharide actively inhibits the addition of a fucose by Fut8 (*58*). This discrepancy may be explained by the alternative binding specificities of certain lectins. E-PHA predominantly binds to N-glycans containing a bisecting structure but has also been shown to have affinity for oligosaccharides with terminal galactose residue (*48*, *59*). Similarly, LCA has very high binding affinity for α1,6-fucosylated glycans, but can also bind the mannose core of an N-glycan in the absence of 213 core fucosylation (*47*). Further investigation will be required to decipher which structures they possess as data from lectin binding suggests the presence of either a core fucosylated N-glycan with a possible terminal galactose residue, or a bisecting N-glycan. Data from previous studies (*53*) along with glycosyltransferase knockdown and chemical inhibition experiments in this study, suggest that at least one of these three sites contains an N-glycan with a bisecting structure, with the most likely candidate being N1587 as mutation of this residue resulted in the greatest reduction in E-PHA binding. Future studies using additional mutations on both the essential asparagine residue and on other proximal non-essential residues could reveal the impact of the amino acid alteration on protein expression, function, and stability. Many studies which involve the mutation of N-glycosylation sites on a glycoprotein also use simultaneous multi-site mutation to investigate whether the combined impact of losing two or more N-glycans differs from the effect observed following loss of a single N-glycan (*52*, *60*, *61*). This is a strategy that could reveal crucial information about the synergistic roles of N-glycans on the Notch1 ECD.

Regarding the mutations at sites N888, N1241 and N1587, their effect specifically prevented canonical Notch signalling. Indeed, lectin binding data suggests that the N-glycan at N888 possessed either a bisecting or core fucosylated structure and in view of the evidence from glycosyltransferase knockdown experiments, it is likely that its absence does not lead to accumulation of Notch1 in the ER-Golgi complex. Furthermore, since the mutation at this site did not result in any change to Notch1 protein levels in the cell, i.e., did not accelerate Notch1 degradation, this suggests that it is most likely a core fucosylated N-glycan (possibly with a terminal galactose residue), rather than a bisecting N-glycan. Taken together, these data suggest that the N-glycan at N888 has a direct influence on Notch1 signal activation at the membrane. This may occur through regulation of processes such as receptor folding or cleavage, mechanisms which are regularly affected by loss of an individual N-glycan on other membrane-bound proteins (*62–64*). The N1241I mutation resulted in a dramatic decrease in Notch1 protein levels in the cell, implying that the Notch1 receptor was rapidly degraded, and as a result, Notch target gene expression was inhibited. While lectin binding data suggests that there may have been no N-glycan present at N1241, there is a strong evidence from the levels of the mutant receptor and downstream signalling and that an N-glycan does indeed reside at this site but does not contain bisecting or core fucosylated structures. Removal of the N-glycan at N1587 also reduced Notch1 protein levels, most likely through receptor degradation. Combined with lectin binding data and evidence from the knockdown studies, the cumulative data suggests that the oligosaccharide which resides at this site contains a complex bisecting structure (Table II).

**Table II:**
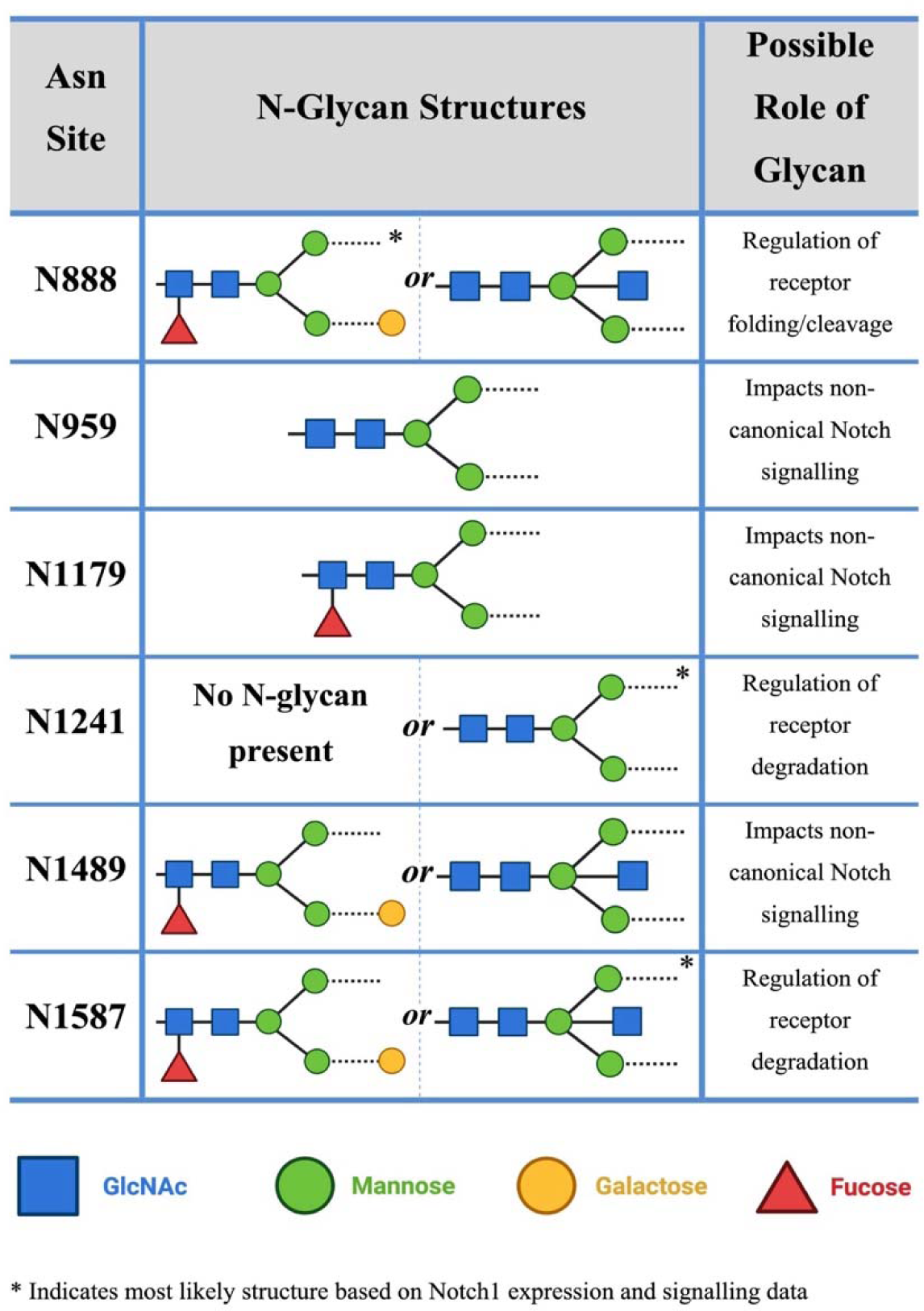
N-Glycan Structures and Possible Roles on the Notch1 Receptor. The N-glycan structures and possible role of each glycan in regulating Notch signalling at specific asparagine residues on the Notch 1 ECD.

Notably, all six mutations inhibited myogenic differentiation of stem cells in response to Jagged-1 stimulation. This further supports lectin binding data by confirming that N-glycans reside at each of the six consensus sites and reinforces data from chemical inhibition and glycosyltransferase knockdown experiments by demonstrating that all these glycans are involved in the process of Notch-mediated myogenic differentiation. However, only the mutations at sites N888, N1241 and N1587 significantly attenuated the expression of the Notch target gene *Hey1*, suggesting that the other three mutations at sites N959, N1179 and N1489 have an indirect effect on myogenic differentiation which is independent of canonical Notch signalling. In addition, none of these three mutations led to significant changes in Notch1 receptor protein levels in the cell. Their effect on myogenic differentiation may be related to non-canonical Notch signalling pathways, such as the Wnt/β-catenin pathway(*65*, *66*) as the Notch1 receptor has been shown to interact with activated β-catenin at the membrane (*67*). While this interaction occurs at the intracellular C-terminal of the Notch1 receptor and it is unlikely that any direct interaction takes place between these glycans and β-catenin, it is possible that their absence leads to altered processing/folding of the receptor which results in dysregulation of Wnt signalling and myogenic differentiation.

Canonical Notch signalling operates via the NICD-RBP-Jκ complex which, along with other transcription factors, activates Notch target gene expression (*2*) to maintain SMC differentiation (*68*, *69*) through interaction with the serum response factor (SRF)-myocardin transcription activator complex (*70*). Several studies have also described the ability of Notch target genes, such as *Hey1* and *Hey2*, to inhibit myocardin-induced SMC differentiation, although many of these studies differ in their description of the mechanism of inhibition, with some citing a direct interaction between Notch target genes and SRF, while others suggest that this occurs via an independent parallel pathway (*71–73*). In addition, CBF1 (the human form of RBP-Jκ) interacts with SMAD2/3 in the TGF-β signalling pathway to promote transcription of SMC genes independent of myocardin, such as *Cnn1* and *SM22*α, albeit this interaction was not observed with Notch1. Notwithstanding the fact that Notch1 NICD-induced signalling in S100β mVSCs promotes Notch target gene and myogenic gene expression, the Jagged-1 response was dependent on canonical Notch1 signalling as the γ-secretase inhibitor, DAPT effectively abrogated this response. In agreement with previous studies, knockdown of *Notch1* mRNA in S100β mVSCs also significantly attenuated the Jagged-1-induced response confirming the pivotal role of Notch1 in lineage specification of these cells *in vitro* and during lesion fromation *in vivo* (*10*, *11*). Future studies that selectively delete the Notch1 receptor in resident S100β mVSCs using *S100*β*-CreERT2* transgenic mice crossed with Notch1 floxed mice before injury or atherosclerotic lesion formation will further confirm the importance of this data *in vivo,* in addition with complimentary studies that explore the role of the other three Notch receptors in this process. In this context, loss of Notch 2 in SMCs leads to protein profile changes during lesion formation but does not alter overall lesion morphology or cell proliferation (*74*). Nevertheless, the current findings highlight the potential of Notch1 as a therapeutic target for controlling the accumulation of SMC-like cells from mVSCs in arteriosclerotic lesion progression (*10*, *11*, *13*)

Limitations of this study include the narrow range of available lectins which are specific for N-linked oligosaccharides. Most commercially available lectins exhibit their highest affinity for a particular monosaccharide in a specific branching/linkage format. However, many of these structures can be found in both O-linked and N-linked glycans, such as Erythrina critagalli lectin (ECL) that bind strongly to the terminal galactose of an N-acetyllactosamine-containing N-glycan, but will also bind to a terminal β1,4-galactose in O-glycans (*75*). In addition, while mutation of specific asparagine residues is a highly specific method, a single change in nucleotide or amino acid sequence can have a dramatic effect on the expression, processing and/or function of the target protein(*61*, *76*). In this study, isoleucine was chosen as the substitute due the ease of mutation (only a single base substitution required for each Asn site) and the non-reactive nature of the side chain. However, the change in side-chain composition may alter the conformation of the receptor or result in issues with steric hindrance (*77*). While glutamine as a substitute is widely used since its structure is the most similar to asparagine, the polar amide group of the glutamine residue may lead to unwanted molecular reactions (*78*). This can be circumvented by mutating the Asn residue to a number of different amino acids, mutating the serine/threonine residue in the consensus sequence, or mutating a proximal amino acid which is not part of the consensus sequence (*79*). Finally, alternative chemical inhibitors such as swainsonine, which blocks α-mannosidase-II activity, could provide greater insights into the role of N-glycans while knockdown of additional glycosyltransferases such as GnT-IV and ST3Gal1 could be useful in deciphering the exact structures of N-linked oligosaccharides on the Notch1 receptor (*80*, *81*).

In conclusion, activation of the Notch1 receptor by its ligand, Jagged-1, stimulates myogenic differentiation of resident S100β mVSCs. Furthermore, N-glycosylation intervention strategies, including chemical inhibition, siRNA knockdown of the GnT-III and Fut8 glycosyltransferases, and site-directed mutagenesis of N-glycosylation sites, reveal that this signalling is heavily regulated by N-glycans which reside at six consensus sequences along the Notch1 ECD. Several key oligosaccharides at these sites were found to contain bisecting or core fucosylated structures. Notch1 expression and trafficking analysis indicated that N-glycans at asparagine (N) 1241 and 1587 protect the receptor from accelerated degradation, while the oligosaccharide at N888 directly affects signal transduction. Conversely, N-linked glycans at the other three sites do not impact canonical Notch1 signalling but still play a role in Notch-mediated myogenic differentiation, possibly via a non-canonical Notch signalling pathway. The role of bisecting and core fucosylated N-glycans in ligand induced activation of the Notch1 receptor may present as a novel therapeutic target for controlling Notch1 signalling and myogenic differentiation of S100β mVSCs during subclinical atherosclerosis where these cells have been reported to play a major role in early lesion development.

## Materials and Methods

### Materials

A full list of the materials is available in Supplementary Materials

### Cell Culture

Cells were grown in tissue culture flasks (25-175 cm^2^) or in 6-well plates in a humidified incubator and were visualised by a Nikon Eclipse TS100 phase contrast microscope. S100β mouse vascular stem cells (mVSCs) were isolated from the aortic arch of female C57BL/6J mice (used in accordance with EU legislation (Directive 2010/63/EU)) as previously described and grown in maintenance medium (MM1) containing DMEM supplemented with 2% chick embryo extract, 1% FBS, 0.02 µg/ml, bFGF basic protein, B-27 Supplement, N-2 supplement, 1% Penicillin-Streptomycin, 50 nM 2-Mercaptoethanol and 100 nM retinoic acid(*10*). These stem cells exhibited a uniform neural-like morphology in low density culture adopting a dendritic-like tree shape and retaining their morphological characteristics at low density throughout repeated passage. Cells were fed with MM1 every 2-3 days and passaged every 3-4 days or when ∼70% confluent. Cells were routinely grown in MM and dissociated with 0.5X TrypLE and expanded. Cells were used at passage numbers below p25. C3H/10T1/2 embryonic fibroblasts have been extensively characterised as multipotent mesenchymal cells (*45*). Clone 8 - CCL-226 cells were grown in EMEM basal media supplemented with 10% heat inactivated FBS and 1% Penicillin Streptomycin. Cells were used at passage numbers below P20 and were always sub-cultured before confluency surpassed 70% to prevent post-confluence inhibition. Freezing media consisted of EMEM with 10% FBS, 1% Pen-Strep and 5% DMSO.

### Adipogenesis Differentiation

Adipogenic differentiation of mVSCs was performed using StemPro® adipogenesis differentiation medium in accordance with the manufacturer’s recommendations. Briefly, cells were seeded at a density of 1 x 10^4^ cells/cm^2^ in their maintenance medium and left for two days before being washed with PBS twice followed by feeding with StemPro® adipogenesis differentiation media for 14 days. Cells were harvested and adipogenic differentiation was evaluated by measuring the expression level of lipoprotein lipase (*Lpl*) and fatty acid binding protein 4 (*Fabp4*).

### Osteogenesis Differentiation

Osteogenic differentiation of mVSCs was performed using the StemPro® osteogenic differentiation medium in accordance with the manufacturer’s recommendations. Briefly, cells were seeded at a density of 1 x 10^4^ cells/cm^2^ in their maintenance medium and left for two days before being washed with PBS twice followed by feeding with StemPro® osteogenic differentiation media for 21 days. Cells were harvested and osteogenic differentiation was evaluated by measuring the expression level of alkaline phosphatase (*Alpl*) and Sclerostin (*Sost)*.

### Telomerase Activity Assay

Cell lysates were prepared using the RIPA buffer method and diluted to a protein concentration of 10-750 ng/μL before telomerase activity was measured using a TRAPeze® XL Telomerase Detection Kit. Briefly, PCR was carried out using fluorescein-labelled primers to amplify telomeric repeats in the sample DNA. Higher telomerase activity meant that there were more telomeric repeats to be amplified by the Taq polymerase, resulting in a more fluorescent product. The PCR products were analysed in a Tecan Safire II fluorescent plate reader. Telomerase activity was calculated by comparing the level of fluorescence detected in each sample to a standard curve generated from a set of telomeric repeat standards. The telomerase activity of a sample was given in TPG units.

### Plasmid Amplification and Purification

#### Transformation

LB Agar (20g/L LB, 12g/L Agar) was made up in dH_2_O and autoclaved to sterilise. After cooling to 55°C, appropriate antibiotics were added (ampicillin at 100μg/mL or kanamycin at 50μg/mL). The agar was poured into sterile petri dishes, allowed to harden, and stored at 4°C. A sample (100ng) of plasmid DNA was mixed with 50μL of competent JM109 bacterial cells in a sterile microfuge tube. Tube was placed on ice for 30 mins. Cells were then heat-shocked at 42°C for 45 secs to allow plasmid DNA to cross the cell membrane. Tube was placed back on ice for 2 mins. 500μL of SOC medium (20% tryptone, 5% yeast extract, 0.5% NaCl, 1% 0.25M KCl, 2% 1M glucose) was added to the tube which was then incubated at 37°C for 45 mins. The contents of the tube were transferred to a LB Agar plate with appropriate antibiotics. Plates were incubated overnight at 37°C. A single colony was picked from the transformation plate and inoculated in 5mL of LB broth (20g/L LB) with appropriate antibiotics. This culture was incubated at 37°C in a shaking incubator for 6-8 hours. 1.2mL of this culture was then transferred to a sterile conical culture flask containing 250mL of LB broth with appropriate antibiotic. Flask was incubated at 37°C overnight in a shaking incubator. 50mL was removed from the flask for plasmid purification. The rest was used to make stocks by supplementing with 50% glycerol and storing at −80°C.

#### Plasmid DNA Purification

Plasmid DNA was purified from transformed bacterial cells using the Invitrogen PureLink^TM^ HiPure Plasmid Filter Midiprep Kit according to the manufacturer’s guidelines. A Nanodrop^®^ ND-1000 Spectrophotometer was used to quantitate the DNA concentration before DNA samples were interrogated by agarose gel electrophoresis and visualised with the SynGene G-Box.

### RNA Extraction

RNA was extracted from cells using the Promega Reliaprep^TM^ RNA Miniprep System according to the provided instructions. A Nanodrop^®^ ND-1000 Spectrophotometer was used to quantitate the RNA concentration before RNA samples were analysed by *Quantitative Reverse-Transcription Polymerase Chain Reaction (qRT-PCR)*

### Quantitative Reverse-Transcription Polymerase Chain Reaction (qRT-PCR)

Quantitative RT-PCR was performed using the Rotor Gene Q from Corbett Research, as previously described (*10*). A ubiquitously expressed reference gene, hypoxanthine guanine phosphoribosyltransferase (*Hprt)* was used as a control. Two different kits were used for this method depending on the primer set: Rotor-Gene SYBR® Green RT-PCR Kit (Qiagen, 204174) and SensiFAST™ SYBR® No-ROX One-Step Kit (BioLine, BIO-72001). Samples were run in triplicate for each primer set.

### Immunocytochemistry

Immunocytochemical analysis of protein expression in cells was performed, as described previously(*10*). Fixed cells on coverslips were stained for expression of the intermediate myogenic differentiation marker Calponin1 (Cnn1) and the calcium binding protein, S100β by incubating overnight at 4°C with antibody diluted to an appropriate concentration with 1% BSA in PBS. Cells were washed x 3 and incubated at room temperature for 1 hr with the secondary antibody (diluted with 1% BSA in PBS), protected from light. Nuclear staining was performed with DAPI which was diluted to 1μg/mL. Coverslips were mounted before cell nuclei and antibody staining was visualised and imaged using the Olympus DP-50 fluorescent microscope. The number of positive cells was counted for each image and divided by the total number of cells in that image to give a ratio, and subsequently a percentage, of antigen-positive cells.

### Transient Transfections

Transfection of both plasmid DNA and single strand siRNA duplexes into cells was performed using the Mirus Bio TransIT-X2® dynamic delivery system, according to the manufacturer’s guidelines after optimization. Cells were grown to ∼ 80% confluency in the maintenance medium before transfection. The transfection mix was made up in a sterile microfuge tube according to kit protocol and incubated at room temperature for 30 minutes to allow transfection complexes to form. This mixture was then added dropwise to the cell culture. Cells were incubated at 37°C for 48 hours before harvesting. Transfection efficiency was assessed using a control GFP-plasmid or TYE 563 DsiRNA (Mirius Bio), a fluorescently tagged siRNA. A non-transfected sample was used as a negative control. Cells were incubated at 37°C for 48 h before being fixed with 3.7% formaldehyde. Nuclei were stained with DAPI, and coverslips were mounted onto slides before being imaged using a Olympus DP-50 fluorescent microscope. The number of cells containing GFP, or dye were counted and graphed as a percentage of the total number of cells. A transfection efficiency of 99.6% was achieved in the dye-transfected sample, compared to 0% in the non-transfected sample.

### Genomic DNA Extraction

Genomic DNA was extracted from cells using the Bioline ISOLATE II Genomic DNA Kit according to the manufacturer’s instructions. Briefly, cells were washed and scraped into PBS and pelleted by centrifugation at 1500 rpm for 5 minutes. The pellet was resuspended in lysis buffer GL. Proteinase K and Lysis Buffer G3 were then added to the sample before incubation at 70°C for 15 minutes. 99% ethanol was added to each sample to adjust DNA binding conditions. Samples were loaded into columns containing a silica membrane and centrifuged at 11,000 x g for 1 minute. The flow-through was discarded and columns were washed twice in wash buffer. Elution buffer was then added to each column. Samples were incubated at room temperature for 1 minute before centrifugation at 11,000 x g for 1 minute. The flow-through contained the purified DNA.

### Protein Isolation and Quantitation

Cells at >80% confluency were scraped into culture media and pelleted by centrifugation. The supernatant was discarded, cells were resuspended in an appropriate volume of RIPA buffer and incubated on ice for 30 minutes to induce cell lysis. The lysate was then clarified by centrifugation at 8,000 x g for 20 minutes at 4°C. The supernatant containing cellular proteins was retained while the cell debris pellet was discarded. The Pierce™ BCA Protein Assay Kit was used to quantitate the concentration of protein in the clarified cell lysate.

### Membrane Fractionation and Protein Isolation

Isolation of proteins from the cell membrane was carried out using the Mem-PER Plus Membrane Protein Extraction Kit according to the manufacturer’s instructions. Cells were harvested and the Mem-PER™ Plus Membrane Protein Extraction Kit was used to perform membrane fractionation on the crude lysate according to the manufacturer’s instructions. Protein concentration in both the cytoplasmic fraction and membrane fraction was quantified by BCA Assay. Both samples were run on an SDS-PAGE precast protein gel and subsequently transferred to a nitrocellulose membrane. Staining with Ponceau S confirmed the difference in the presence of protein band profiles between the two samples. Immunoblotting was then carried out with an anti-Notch1 antibody and chemiluminescent staining was used to visualise bands.

### Effect of TNC and DMJ on Membrane Localisation of Notch1

Cells were treated in the absence or presence of TNC (0.5μg/mL), DMJ (4μg/mL), 1 x Brefeldin A (BFA) at 37°C for 48 h before being harvested. Membrane fractions were isolated using the Mem-PER™ Plus Membrane Protein Extraction Kit, according to the manufacturer’s instructions. Protein concentration in the membrane fractions was measured by BCA Assay. Western blot was then carried out and samples were immuno-probed using an anti-Notch1 antibody. Chemiluminescent staining was used to visualise immunoblotting. Densitometry was carried out using ImageJ to measure the intensity of Notch1 bands while β-actin, pan-cadherin and Ponceau S staining was used as loading controls.

### Trafficking of Notch1 Receptor

Cells were seeded at 5,000 cells/well onto sterile coverslips and treated with or without TNC (0.5μg/mL), DMJ (4μg/mL), 1 x BFA at 37°C for 48 h before being fixed with 3.7% formaldehyde. Immunocytochemistry was then carried out and cells were stained using an anti-Notch1 antibody with an IgG control antibody used as a secondary control. A control sample was also stained with an anti-Golgin-97 antibody to visualise the subcellular location of the golgi apparatus (GA). Nuclei were stained with DAPI. Samples were imaged using a fluorescent microscope. ImageJ was used to calculate the mean fluorescence intensity and the integrated density of fluorescence at both the cell membrane and in the perinuclear region (adjacent to the nucleus, i.e., where the GA is located). Samples were graphed as a ratio of the fluorescence at the membrane to the fluorescence in the perinuclear region.

### Purification of Notch1 by Immunoprecipitation

Protein lysates were prepared, and the protein concentration measured by BCA assay. Immunoprecipitation was carried out using Protein A-conjugated magnetic beads and either an anti-Notch1 rabbit monoclonal antibody, a rabbit IgG control antibody, or a mouse IgG control antibody. Following incubation with the antibody, supernatants (containing unbound proteins) were transferred to a fresh tube and appropriately stored. Bound protein was eluted off the beads by either 1X Laemmli Buffer or a 0.1M Glycine buffer (pH 2.0). Samples eluted with the glycine buffer were neutralised using a 1M Tris buffer (pH 9.0). Both eluted bound proteins and previously stored unbound protein samples were then run on an SDS-PAGE precast protein gel, transferred to a nitrocellulose membrane, immuno-probed using an anti-Notch1 antibody and visualised by TMB staining of the HRP-conjugated secondary antibody.

### Ectopic Expression of Myc-tagged Notch1 and NICD

The pCS2 Notch1 full length-6MT and pCS2 Notch1 ICv-6MT plasmids were purchased from Addgene (LGC Standards, Teddington, UK) as bacterial stabs. Plasmid-containing bacteria were transferred to a 5mL culture with selective antibiotic at 37°C for 8 hrs. 1.2mL of this culture was then transferred to a 250mL secondary culture with selective antibiotics which was incubated overnight at 37°C. Plasmid DNA was isolated from this culture using the Invitrogen PureLink^TM^ HiPure Plasmid Filter Midiprep Kit. To verify plasmid length and quality, plasmid DNA was digested with an appropriate restriction enzyme. This served to linearise the DNA before running plasmid samples through gel electrophoresis on an agarose gel. Visualisation of this gel under UV light confirmed that plasmids were the correct length and that the stocks contained no contaminating DNA. To validate the ectopic expression of both proteins, cells were grown to ∼80% confluency in 3 wells of a 6-well plate and transfected with either the Notch1 full-length plasmid, the NICD plasmid or an empty vector plasmid (pCDNA3.1) using the TransIT-X2 Dynamic Delivery System. Cells were incubated at 37°C for 48 h in MM1 before being harvested. Protein was isolated and quantified by BCA assay. Western blot was then carried out, with all samples being immuno-probed using an anti-Myc-tag antibody to stain for either the myc-tagged Notch1 or myc-tagged NICD, and an anti-β-actin antibody was used as the loading control.

### Western Blot

Following protein isolation and quantitation, samples were prepared for SDS-PAGE using a precast SDS 10% polyacrylamide gel to perform electrophoresis and the Pierce™ G2 Fast Blotter for protein transfer, as previously described (*68*). Ponceau S staining confirmed that the transfer had been successful. Blocking buffer (5% dried milk in TBS with 0.1% Tween) was added to the membrane and incubated at room temperature for 1-2 h on a rocker before the membrane was washed in TBST (50mM Tris, 150mM NaCl, 1mM MnCl_2_, 1mM CaCl_2_, 1mM MgCl_2_, 0.1% Tween-20) and incubated with the primary antibody (diluted with 5% BSA in TBST) overnight on a rocker at 4°C. Primary antibody was removed, membrane washed 3 times (5 minutes each) with TBST, before incubation with an HRP-conjugated secondary antibody (diluted with 5% BSA in TBST) on a rocker for 1 hr at room temperature. Fresh TMB was added to the membrane which was then incubated on a rocker at room temperature for approx. 10 mins before the blot was imaged using BioRad GelDoc EZ Gel System. Alternatively, Pierce™ ECL Western Blotting Substrate was added to the membrane which was then incubated on a rocker at room temperature for 5 minutes. The blot was then imaged using Syngene G-Box.

### Enzyme-Linked Lectin Assay (ELLA)

All samples were diluted to the same concentration with 50μL of treated sample or control sample added in triplicate to each well of a 96-well plate. The plate was incubated overnight at 4°C with gentle shaking. Wells were emptied and blocking buffer (0.5% PVA in PBS) added to each well. The plate was incubated for 2 hrs at room temperature with gentle shaking before wells were washed with TBST (50mM Tris, 150mM NaCl, 1mM MnCl_2_, 1mM CaCl_2_, 1mM MgCl_2_, 0.1% Tween-20). Biotinylated lectins (50μL, 5μg/mL) were added to each well and incubated at room temperature for 1 hr with gentle shaking before wells were emptied and washed again with TBST. HRP-conjugated anti-biotin antibody (50μL, 1:10,000) was then added to each well and incubated at room temperature for 1 hour with gentle shaking. Wells were washed with TBST before a TMB substrate solution (90μL) was added and incubated for 10 minutes at room temperature with gentle shaking. Stop solution (10% H_2_SO_4_) was added to each well before the absorbance was measured at 450 nm in a plate reader.

### N-glycosylation Profile of Purified Notch1

Cells were grown to ∼80% confluency in 3 wells per 6-well plate. One well was not transfected while the TransIT-X2 Dynamic Delivery System was used to transfect the other two wells with either the Notch1 full-length plasmid or an empty vector control plasmid. Cells were incubated at 37°C for 48 h in maintenance media (MM1) before being harvested. Protein was isolated and quantified by BCA assay. IP was carried out to purify the ectopically expressed Notch1 using Protein G-conjugated magnetic beads and an anti-Myc-tag antibody. The non-transfected sample was immunoprecipitated with a mouse IgG control antibody. Bound proteins were eluted off the magnetic beads using a 0.1M Glycine buffer (pH 2.0) and a 1M Tris buffer (pH 9.0) was used to neutralise the purified protein fraction. BCA assay was carried out again to determine the concentration of protein in each sample post-IP. ELLA was then applied to detect the presence of N-glycans in all three samples. Control wells containing PBS rather than a protein sample were used to determine the baseline for non-specific absorbance, while the mouse IgG control sample acted as the baseline for non-specific lectin- or antibody-binding to the IP eluate. Both baseline values were subtracted from sample absorbance values to decipher the true level of specific lectin binding. TBST acted as the vehicle control for all lectins.

### Site-directed mutagenesis

Mutation of N-glycosylation sites on a plasmid-encoded Notch1 receptor was achieved using the Agilent QuikChange Lightning Site Directed Mutagenesis Kit, according to the manufacturer’s instructions. Briefly, primers were specific for a region of the Notch1 gene containing an N-glycosylation site and were designed to incorporate a single base substitution in copies of the plasmid DNA template. PCR tubes were placed in an Applied Biosystems® Veriti® Thermal Cycler. A DpnI restriction enzyme was added to each tube of PCR products to degrade any parental template DNA. Mutant plasmids were then transformed into ultracompetent XL10-Gold bacterial cells via heat-shock at 42°C. After allowing the cells to recover for 1 hour in NZY^+^ broth at 37°C, each transformation reaction was spread on an LB agar plate with Ampicillin (50μg/mL) and incubated at 37°C overnight. Colonies were picked and inoculated in liquid LB. Transformed bacteria were grown and mutant plasmids were purified before sending them for sequencing with Source Biosciences to confirm that the correct single base substitution had been incorporated in each case.

### Data Analysis and Statistics

The mean and standard deviation (or standard error margin) was calculated for all the technical and biological replicates in each experiment and these values were used to plot the data. The Kolmogorov-Smirnov normality test was used to analyse the spread of the data as it uses the most stringent conditions. The outcome of the normality test (along with the number of samples and parameters) dictated what type of significance test was carried out. In the case of data containing only two samples, an unpaired t-test with Welch’s correction was performed on normally distributed data. For non-parametric (skewed) data, the Wilcoxon test was carried out to calculate statistical significance. For data containing more than two samples, ANOVA was used to perform multiple comparisons between all samples with an ordinary one-way (one parameter to compare) or two-way (two parameters to compare) ANOVA for normalised data, or the Kruskal-Wallis test for non-parametric data.

## Funding

Irish Research Council (IRC) GOIPG/2016/1529 (EC)

Irish Research Council (IRC) GOIPG/2014/43 (MDL)

Government of Saudi Arabia ID: IR15208 (JG)

National Institutes of Health R01-AA024082 (EMR)

Science Foundation Ireland (SFI) award 11-PI-1128 (PAC)

Health Research Board (HRB), Ireland HRA-POR-2015-1315 (PAC)

## Author contributions

Conceptualization: PAC, EMR, EC

Methodology: EC, AO, BOC, YG, RH, MDL

Investigation: EC, AO, MDL, YG, RH

Visualization: EC, PAC

Supervision: PAC

Writing—original draft: EC, AO

Writing—review & editing: AO, EMR, PAC

## Data and materials availability

The datasets used and/or analysed during the current study are available from the corresponding author on reasonable request. All data are available in the main text or the supplementary materials.

---

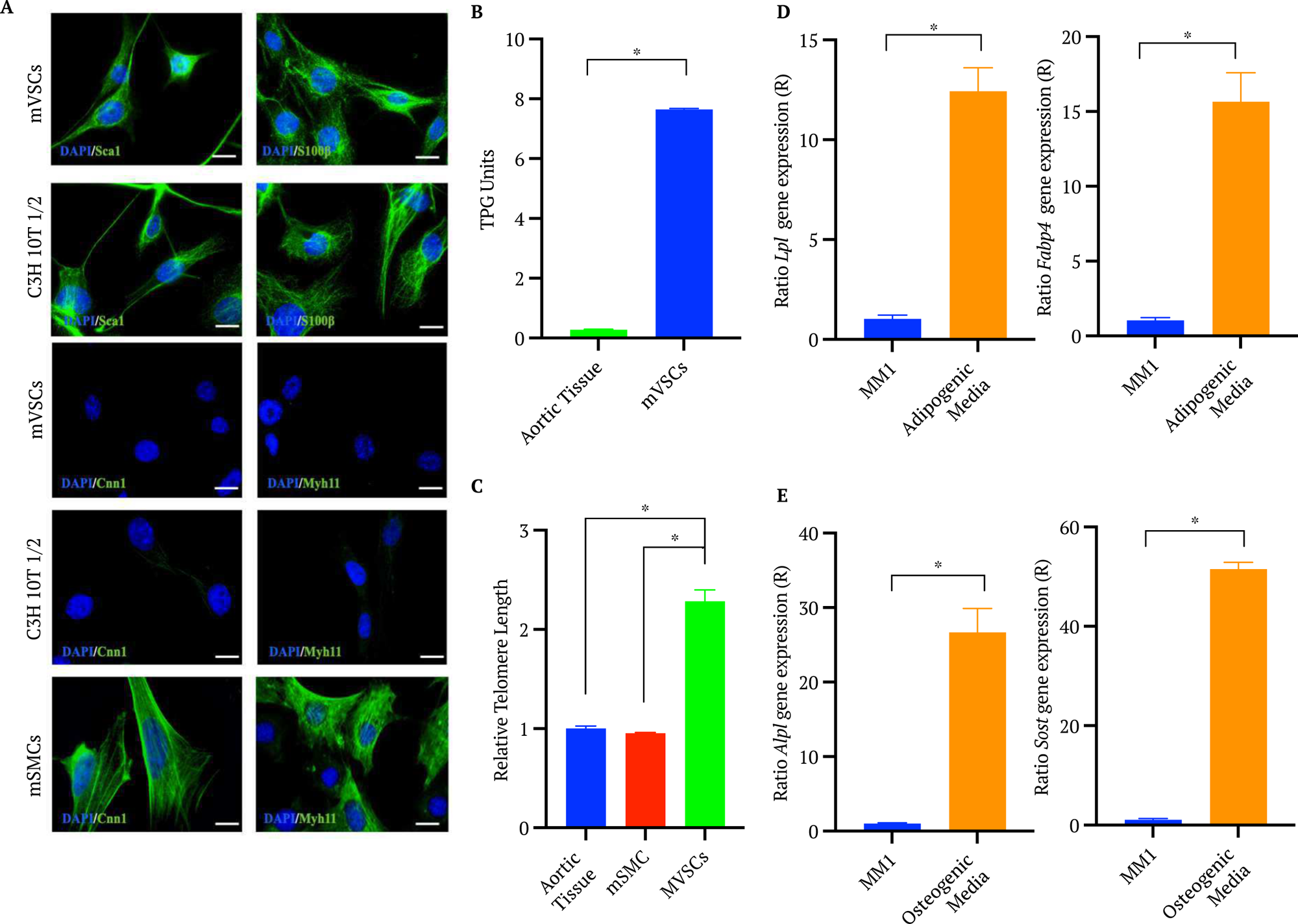

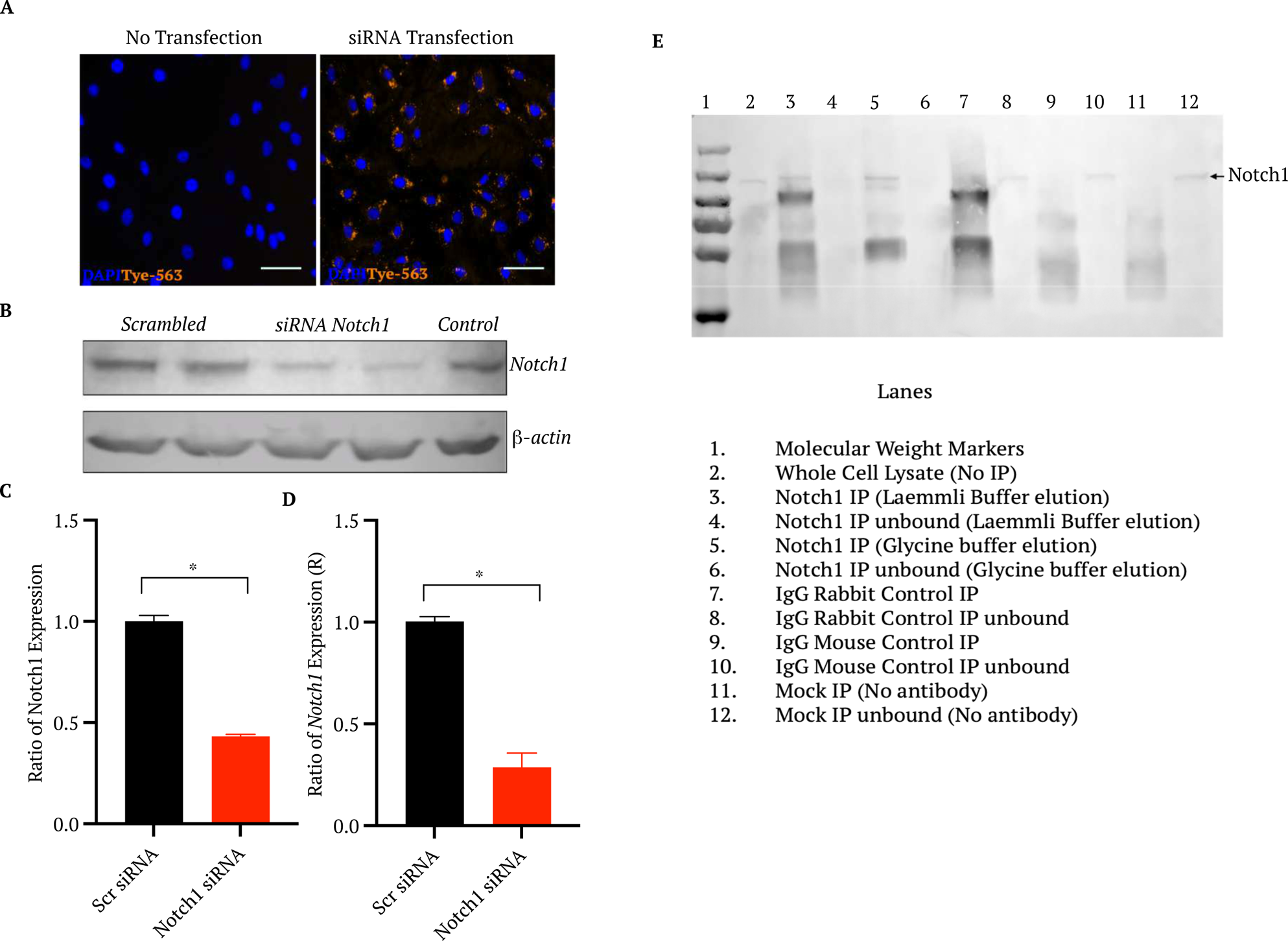

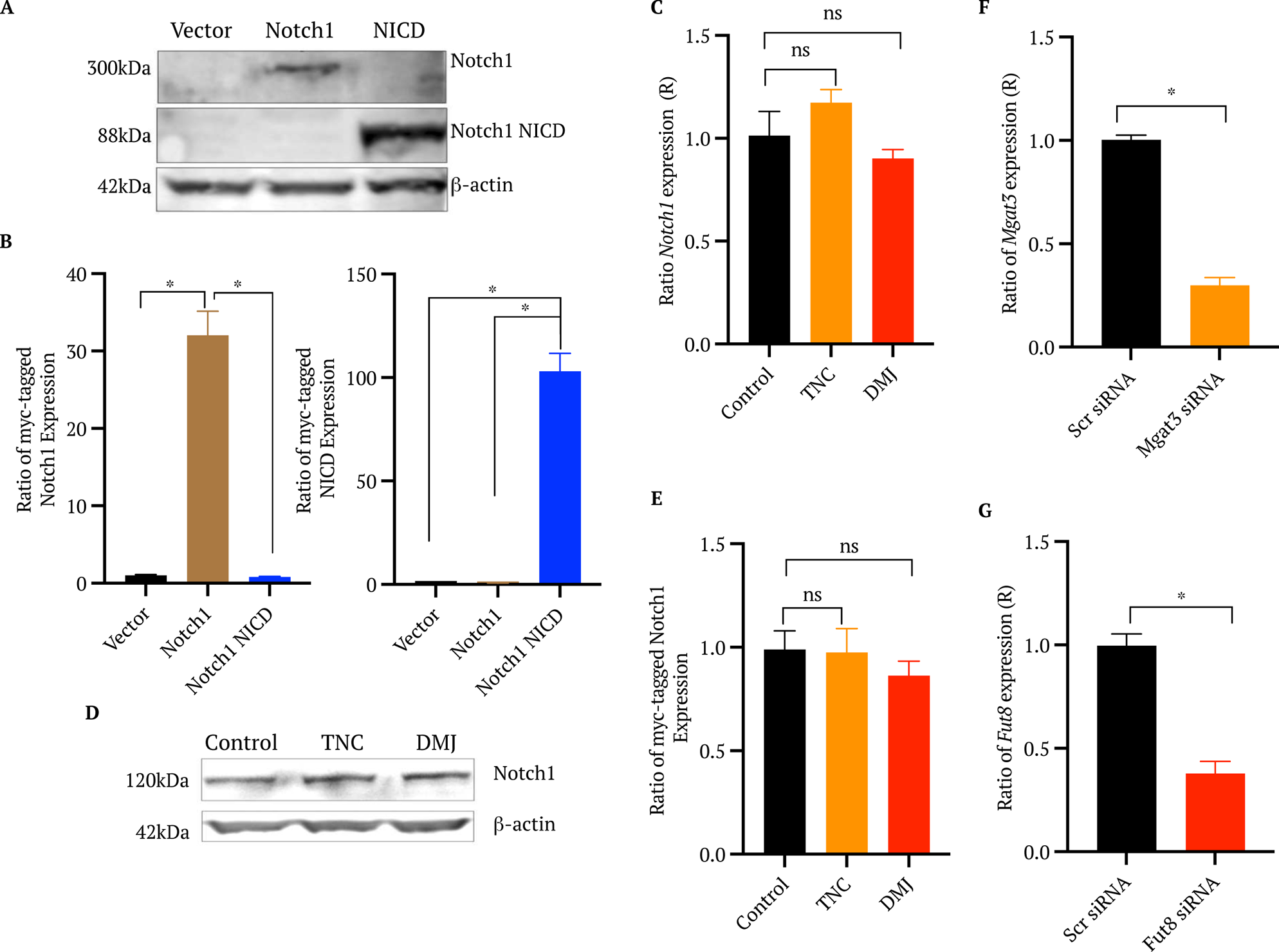

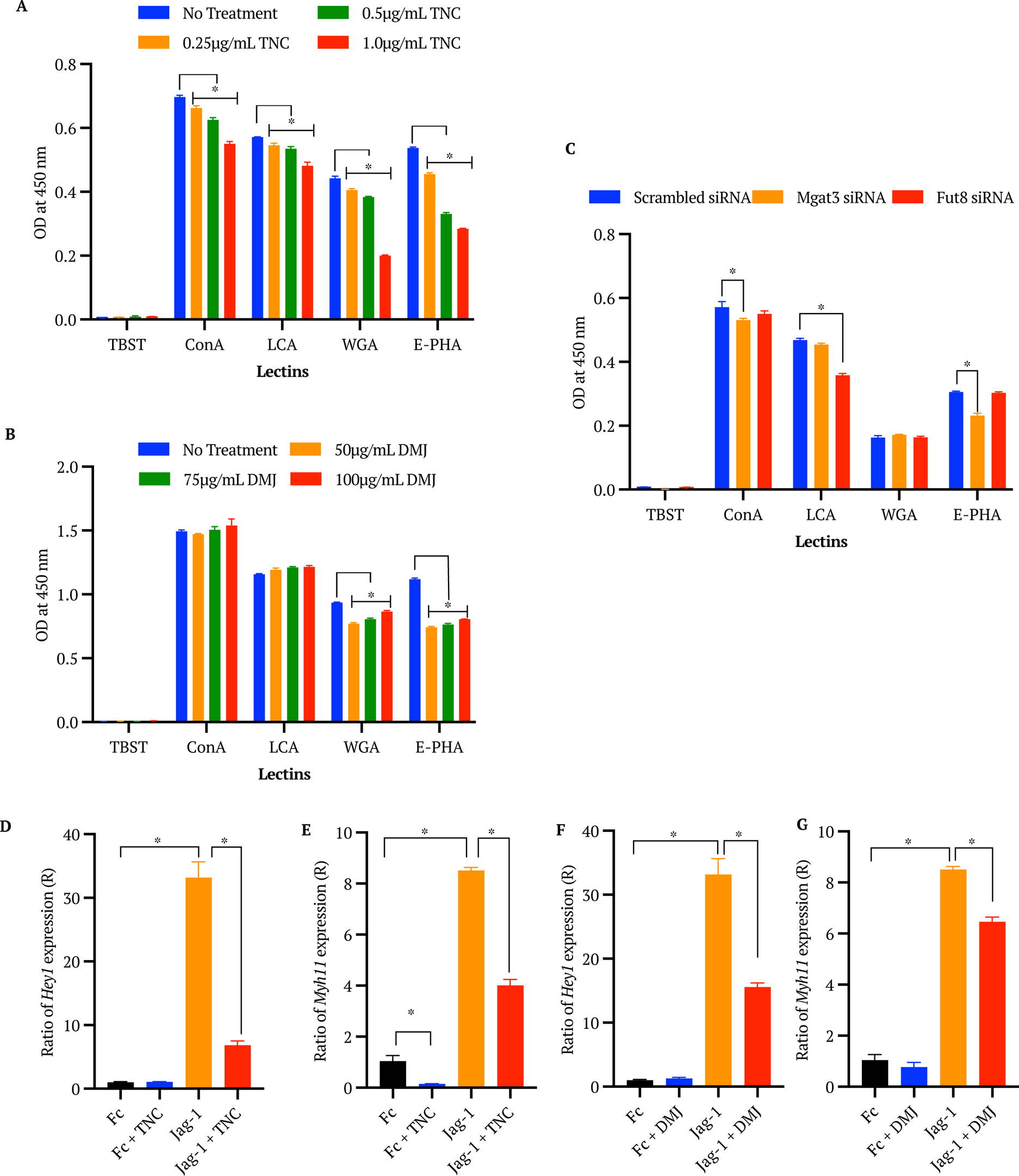

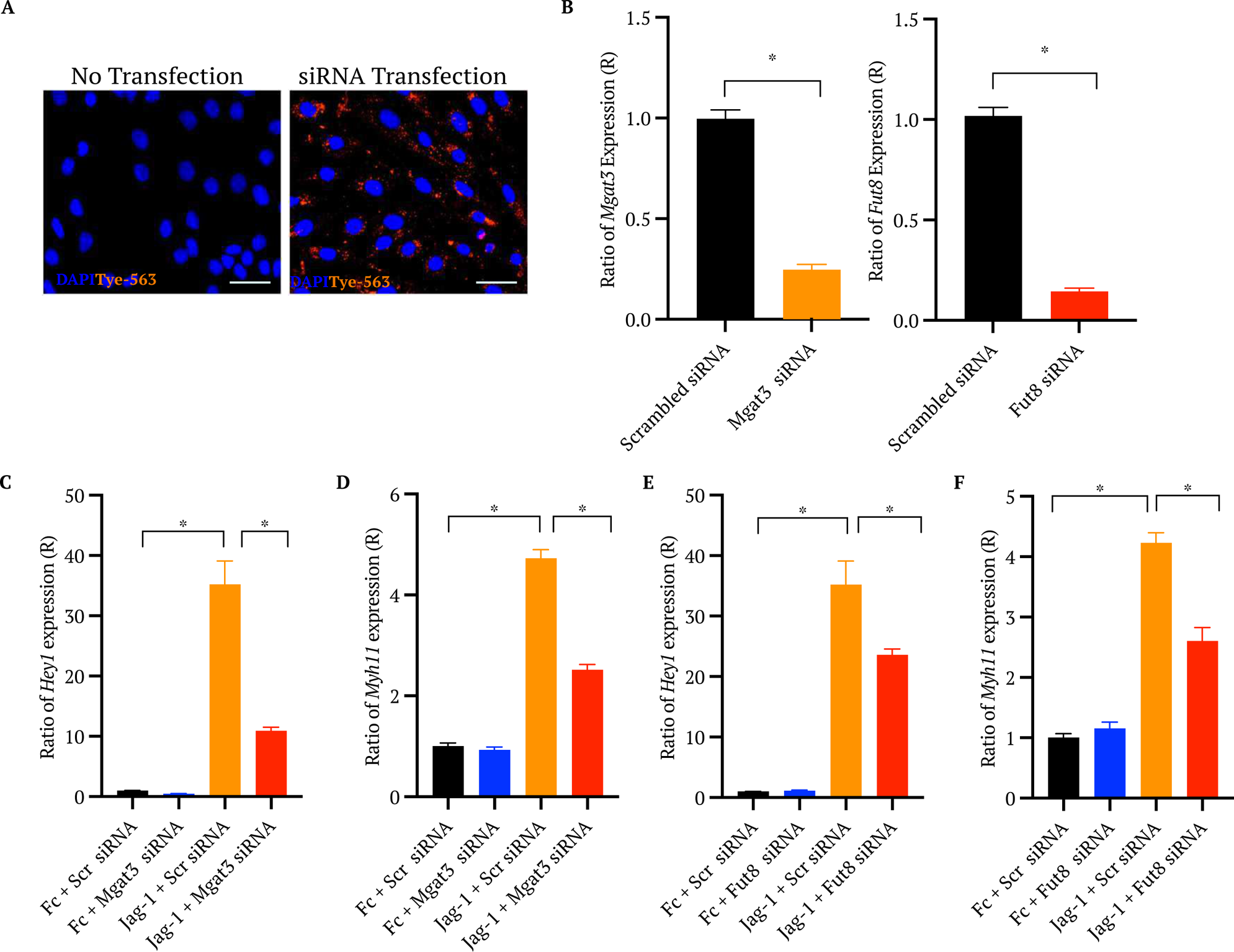

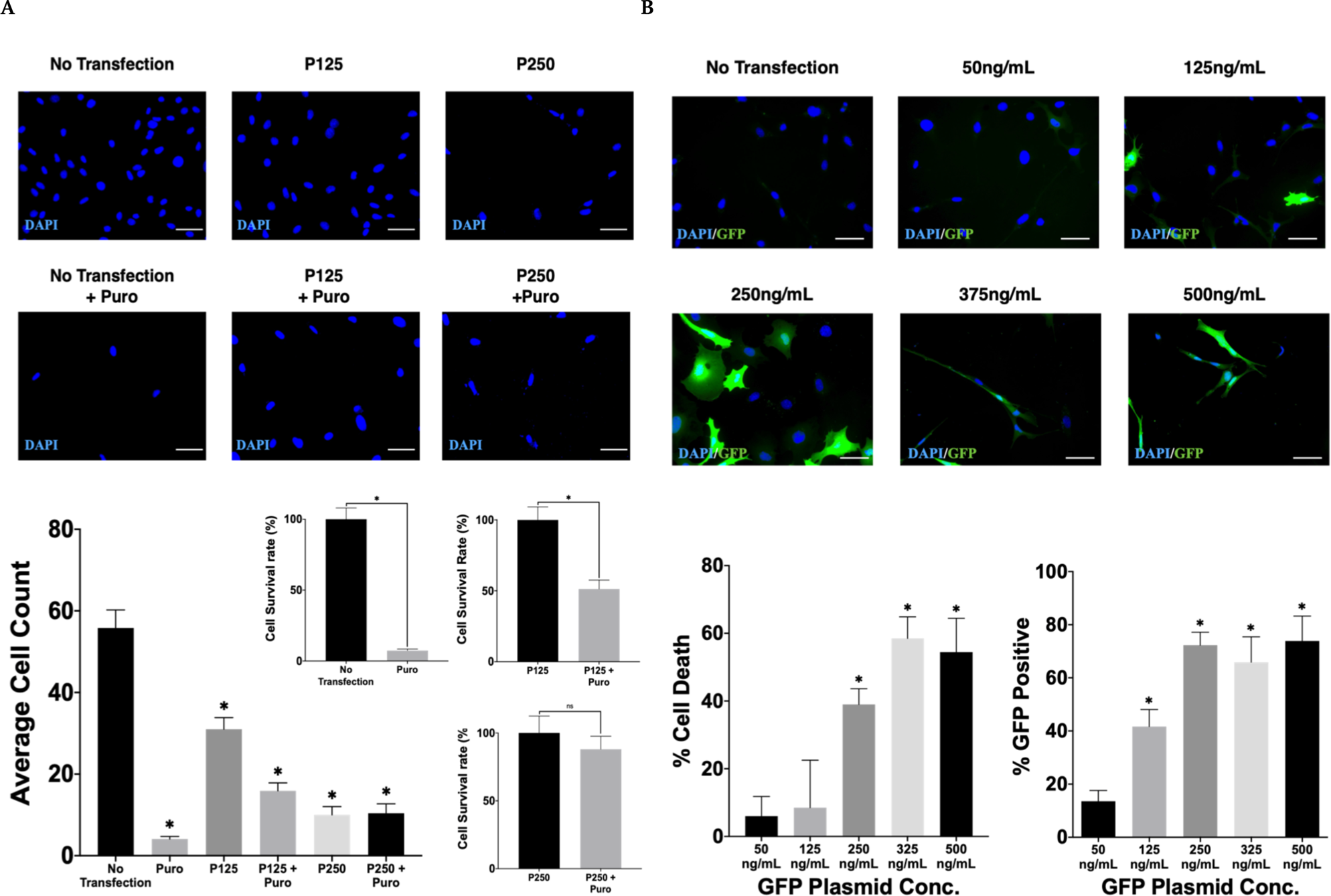

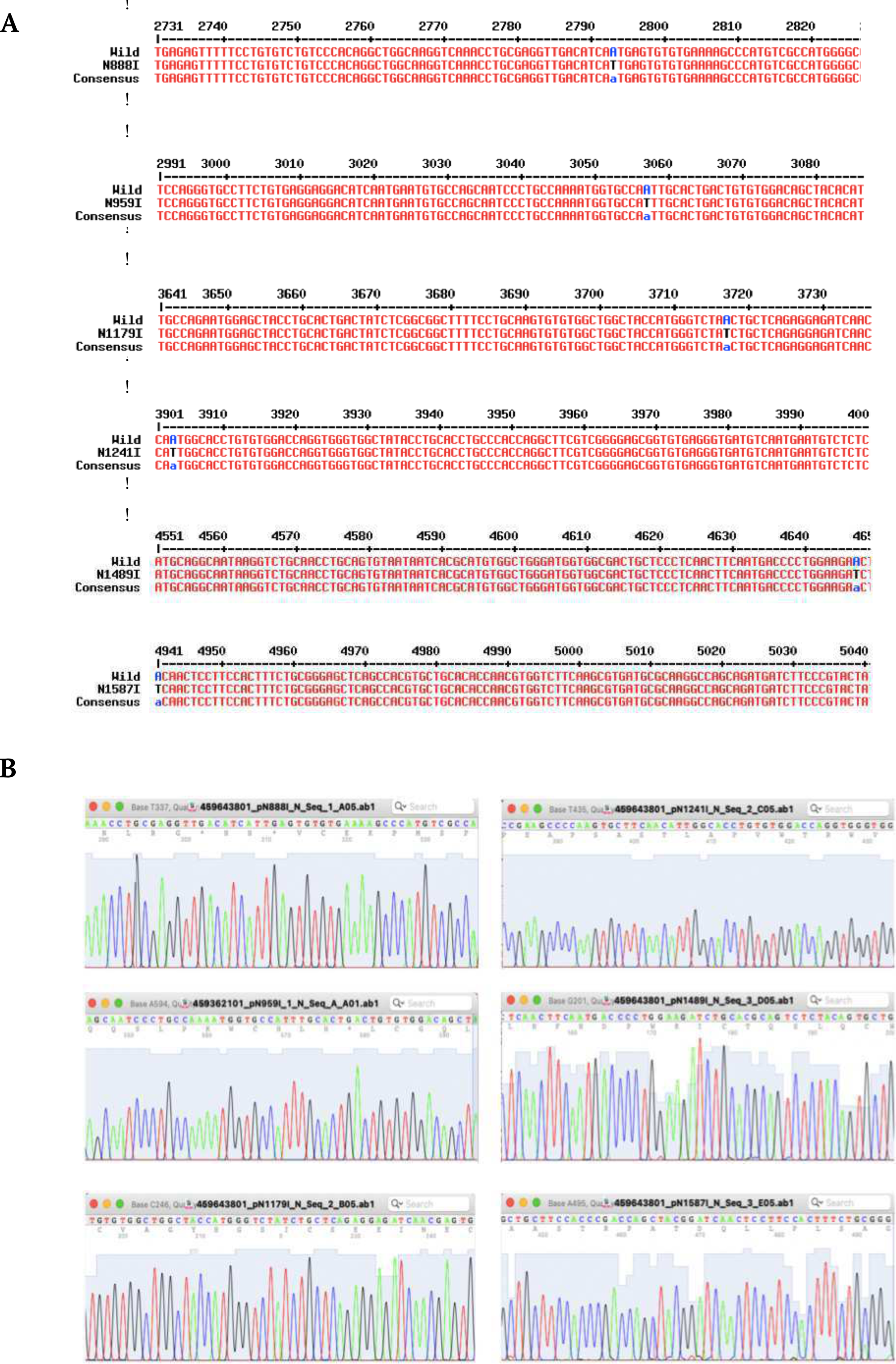

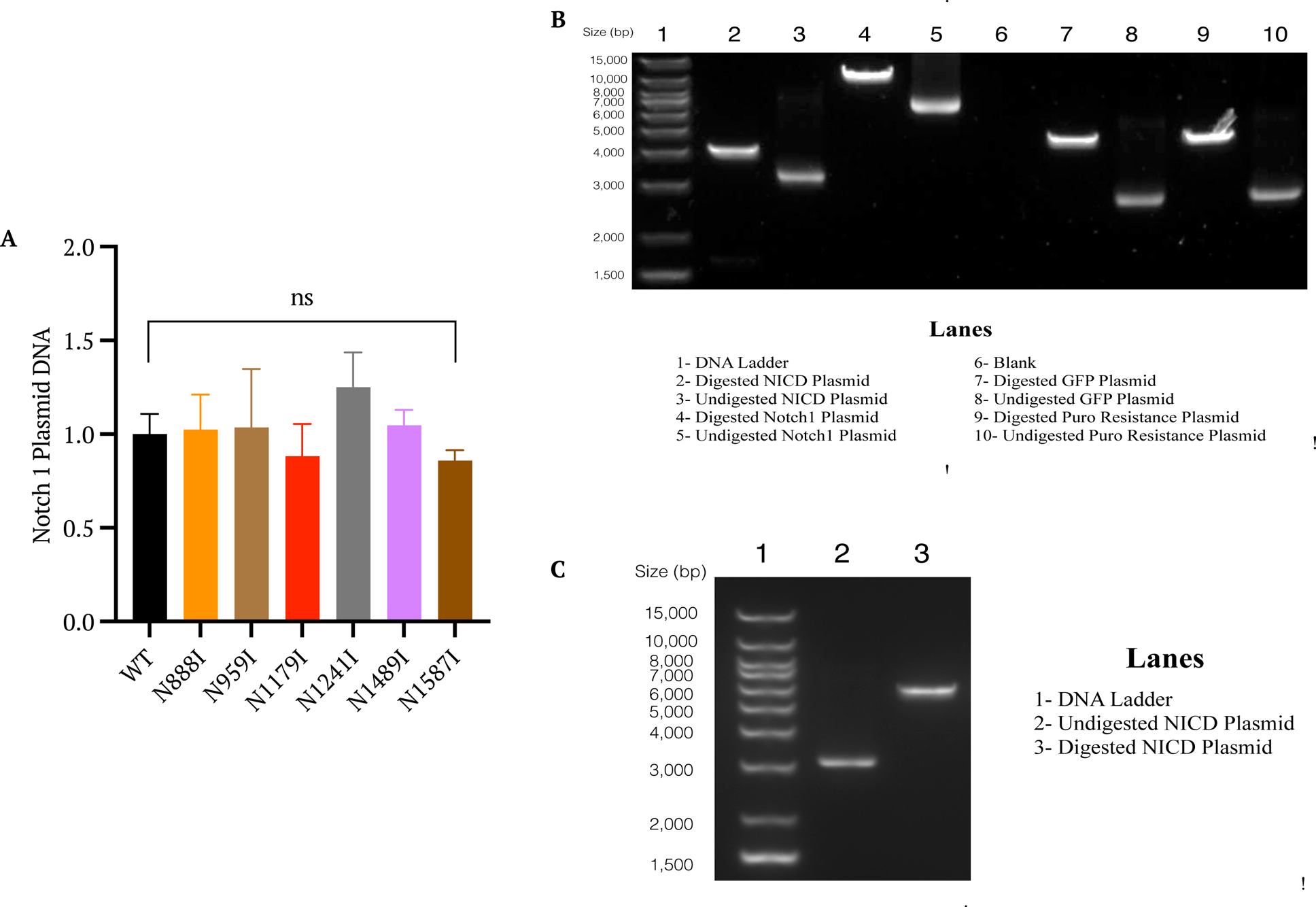

