## Supplemental Figures and Materials for "N-Glycans on the extracellular domain of the Notch1 receptor control Jagged-1 induced Notch signalling and myogenic differentiation of S100β resident vascular stem cells"

Eoin Corcoran *et al.*

Supplementary Text

**Mammalian Cell Lines**

| **Cell Line** | **Cell Type** | **Description** |
| --- | --- | --- |
| mVSC | Adult Stem Cell | Multipotent Vascular Stem Cells isolated |
|  |  | from the medial layer of a mouse aortic |
|  |  | arch by enzymatic dispersion and |
|  |  | sequential plating on non-adherent and |
|  |  | adherent tissue culture plates |
| MOVAS (ATCC® | Differentiated | Commercial cell line of terminally |
| CRL-2797™) | Smooth Muscle Cell | differentiated mouse aortic smooth |
|  |  | muscle cells. Immortalised with SV40 |
|  |  | large T-antigen |
| C3H/10T1/2, Clone | Embryonic fibroblast | Commercial cell line of fibroblasts |
| 8 (ATCC® CCL- |  | isolated from the embryo of C3H mice. |
| 226™) |  |  |

**Bacterial Cell Lines**

**Cell Line	Cell Type	Genotype**

*E.coli* JM109	Bacterial Cell	endA1, recA1, gyrA96, thi, hsdR17 (rk–,

mk+), relA1, supE44, Δ( lac-proAB), [F´ traD36, proAB, laqIqZΔM15]

XL-10 Gold	Ultracompetent Bacterial Cell

Tetr∆ (mcrA)183 ∆(mcrCB-hsdSMR- mrr)173 endA1 supE44 thi-1 recA1 gyrA96 relA1 lac The [F’ proAB lacIqZ∆M15 Tn10 (Tetr) Amy Camr]

**Primers Qiagen**

| **Product** | **Target Gene** | **Catalog No.** |
| --- | --- | --- |
| Mm_Hprt_1_SG QuantiTect Primer Assay | *Hprt* | QT00166768 |
| Mm_*Mgat3*_1_SG QuantiTect Primer Assay | *Mgat3* | QT00297073 |
| Mm_*Fut8*_3_SG QuantiTect Primer Assay | *Fut8* | QT00115094 |

**Primers Integrated DNA Technologies**

**Target Gene	Sequence**

*Hprt*

Forward 5'- GGC TAT AAG TTC TTT GCT GAC CTG C -3'

Reverse 5'- GCT TGC AAC CTT AAC CAT TTT GGG -3'

*Hey1*	Forward 5'- ACT CCG ATA GTC CAT AGC CA -3' Reverse 5'- GTA CCC AGT GCC TTT GAG AA -3'

*Cnn1*	Forward 5'- TCC ATG AAG TTG TTC CCG ATG -3'

Reverse 5'- GCT TGT CTG CTG AAG TAA AGA AC -3' Forward 5'- ATT GTC ATT TAG CGG GTC CAT -3

*Myh11*

*Notch1*

Reverse 5'- AAC AGA GTT CTC CAT CAT CCA C -3' Forward 5’ – AGG ATC AGT GGA GTT GTG C – 3’

Reverse 5’ – CGT TAC ATG CAG CAG TTT CTG – 3’

**Plasmid Vectors**

**Plasmid	Description**

pCS2 Notch1 Full Length- 6MT

pCS2 Notch1 ICv-6MT

Plasmid encoding for the full-length mouse Notch1 protein with a hexameric myc-tag at the C-terminus end. Prof.

Raphael Kopan (Addgene plasmid No. 41728)

Plasmid encoding for the intracellular domain of mouse Notch1 protein (aa1744 – 2184) with a hexameric myc-tag at the C-terminus end. Prof. Raphael Kopan (Addgene plasmid No. 41730)

pEGFP-N1	Plasmid encoding for Enhanced Green Fluorescent Protein.

Dr. Ronan Murphy, Dublin City University, Ireland.

pPGKpuro	Plasmid encoding a Puromycin Resistance gene. Dr. Dermot Walls, Dublin City University, Ireland

pCDNA3.1	Empty plasmid vector containing only bacterial resistance genes and restriction sites (Addgene plasmid no. V790-20)

**siRNA Duplexes**

**Product	Target Gene	Catalog No.**

Silencer® Select siRNA, Notch1 (s70700)	*Notch1*	4390771 Silencer® Select siRNA, *Mgat3* (s69837)	*Mgat3*	^4390771^

Silencer® Select siRNA, *Fut8* (s79249)	*Fut8*	4390771 Silencer® Select GAPDH Positive Control siRNA *Gapdh*	4390849 Silencer® Select Negative Control No. 1 siRNA	No Target	4390843 Silencer® Select Negative Control No. 2 siRNA	No Target	^4390846^

IDT TYE 563 Transfection Control DsiRNA	No Target	51-01-20-19

**Antibodies**

| **Supplier** | **Product** | **Description** | **Catalog No.** |
| --- | --- | --- | --- |
| Abcam | Anti-Calponin Antibody | Rabbit monoclonal | ab46794 |
|  | (EP798Y) |  |  |
|  | Anti-smooth muscle Myosin | Mouse monoclonal | ab683 |
|  | heavy chain 11 antibody |  |  |
|  | (1G12) |  |  |
|  | Anti-myc-tag Antibody | Mouse monoclonal | ab32 |
|  | (9E10) |  |  |
|  | Anti-S100β Antibody | Rabbit monoclonal | ab52642 |
|  | (EP1576Y) |  |  |
|  | Anti-Nestin Antibody | Mouse monoclonal | ab6320 |
|  | (196908) |  |  |
|  | Anti-pan Cadherin Antibody | Rabbit monoclonal | ab51034 |
|  | [EPR1792Y] |  |  |
|  | Anti-Notch1 Antibody | Rabbit monoclonal | ab52627 |
|  | [EP1238Y] |  |  |
| Biosciences | Anti-Golgin-97 Antibody | Mouse Polyclonal | PA530048 |
|  | Anti-Notch1 Antibody (A6) | Mouse Monoclonal | MA511961 |
|  | AlexaFluor® 488 Anti-Rabbit IgG | Goat polyclonal secondary antibody | A11008 |
|  | AlexaFluor ® 488 Anti- Mouse IgG | Goat polyclonal secondary antibody | A11001 |

| Brennan & Co | Anti-Notch1 Antibody (D1E11) XP® | Rabbit Monoclonal | 3608S |
| --- | --- | --- | --- |
| Sigma- | Anti-β-actin Antibody (AC- | Mouse Monoclonal | A2228 |
| Alrich | 74) |  |  |
|  | HRP-conjugated Anti-Mouse | Goat polyclonal | A9917-1ML |
|  | IgG (Fab specific) | secondary antibody |  |
|  | HRP-conjugated Anti-Rabbit | Goat polyclonal | A0545-1ML |
|  | IgG (whole molecule) | secondary antibody |  |
|  | HRP-conjugated Anti-biotin | Goat polyclonal | A4541 |
|  | IgG |  |  |

**Chemicals, Reagents, Proteins and Kits**

| **Product** | **Supplier** | **Catalog No.** |
| --- | --- | --- |
| Abcam | TEO-Tricine Precast Gels –  RunBlue™ 10% | ab119202 |
| Accuscience | Bio-Rad SureBeads™ Protein G Magnetic Beads | 1614023 |
|  | Bio-Rad SureBeads™ Protein A Magnetic Beads | 1614013 |
| Agilent | QuikChange Lightning Site Directed Mutagenesis Kit | 210518 |
| Biosciences | TrypLE Select | A1217701 |
|  | Opti-MEM | 11058021 |
|  | N2 | A1370701 |
|  | B27 | 17504-044 |
|  | 2-mercaptoethanol | 31350-010 |
|  | UltraPure™ 0.5M EDTA, pH 8.0 (4 x 100ml per box) | 15575020 |
|  | Pierce™ BCA Protein Assay Kit | 23227 |
|  | Invitrogen PureLink™ HiPure Plasmid Filter | K210014 |
|  | Pierce™ ECL Western Blotting Substrate | 32209 |
|  | 1kb Plus DNA Ladder | 10787018 |
|  | Mem-PER™ Plus Membrane Protein | 89842 |

| Bio-Techne | Recombinant Rat Jagged-1 Fc | 599-JG-100 |
| --- | --- | --- |
|  | Recombinant Human IgG1 | 110-HG-100 |
| Fisher Scientific | Day Impex™ Disinfectant Virkon Virucidal Tablets | 12328667 |
|  | PBS Tablets | 12821680 |
| Fluka | Calcium Chloride Solution | 21114-1L |
| Lennox | Industrial Methylated Spirits | CRTSI0330716 |
| LGC Standards | Fetal Bovine Serum, ES Qualified | ATCC SCRR-30-2020 |
| Medical Supply Company | TransIT-X2® Dynamic Delivery System 0.3ml | MIR6003 |
|  | SensiMix SYBR No-ROX Kit (500 rxn) | QT650-02 |
|  | SensiFAST™ SYBR^®^ No- ROX One-Step Kit | BIO-72005 |
|  | Bioline ISOLATE II Genomic DNA Kit | BIO-52067 |
|  | Protein G | 21193-5mg |
| MyBio | Promega Reliaprep RNA cell mini prep system | Z6012 |
| Qiagen | Rotor-Gene SYBR Green RT-PCR Kit (400) | 204174 |
| ScienCell | Relative Mouse Telomere length quantification qPCR | SC-M8908 |
| Seralab | Chick Embryo Extract | CE-650-J |
| Sigma-Aldrich | DAPI | D9542-5MG |
|  | Formaldehyde Solution | 252549-100ML |
|  | Polyvinyl Alcohol | P8136-250g |
|  | Trizma base | T1503-1kg |
|  | Sodium Chloride | S3014-5KG |
|  | Hydrochloric Acid | H1758-500ML |
|  | Manganese(II) Chloride Solution | M1787-100ML |

| Sigma-Aldrich (contd.) | DPBS | D5773-1L |
| --- | --- | --- |
|  | Magnesium Chloride | M1028-100ML |
|  | Solution |  |
|  | D-(+) Glucose | 47829 |
|  | Magnesium Sulfate | M7506-500G |
|  | Puromycin | P8833-10mg |
|  | Ampicillin sodium salt | A9518-5G |
|  | Trypan Blue | T8154-100ML |
|  | Sodium Citrate | W302600 |
|  | Citric Acid | C2404 |
|  | TMB Tablets | T3405 |
|  | Hydrogen Peroxide | 18312 |
|  | RIPA Buffer | R0278-50ml |
|  | Protease inhibitor cocktail | P8340 |
|  | 2-propanol | I9516 |
|  | Ethanol | E7023-500ML-D |
|  | Penicillin-Streptomycin | P4333-100ML |
|  | Bovine Serum Albumin | A4503-50G |
|  | RPMI | R8758-500ML |
|  | EMEM | M4655-500ML |
|  | Fetal Bovine Serum | F9665-500mL |
|  | DMEM | D5796-500mL |
|  | DAPT | D5942-5MG |
|  | DMSO | 41639-500ML |

| Sigma-Aldrich (contd.) | Glycine | G8898-1kg |
| --- | --- | --- |
|  | Tween-20 | P1379-500ML |
|  | Retinoic Acid | R2656 |
|  | Agar | A1296-100G |
|  | Agarose | A9414-5G |
|  | TE Buffer | 93283-100ML |
|  | TRAPeze® XL Telomerase Detection Kit | S7707 |
|  | Triton X-100 | T8787 |
|  | Fluoromount Aqueous Mounting Medium | F4680 |
| Vector Laboratories | Concanavalin A | B1005 |
|  | Lens Culinaris Agglutinin | B1045 |
|  | Wheat Germ Agglutinin | B1025 |
|  | Phaseolus Vulgaris  Erythroagglutinin | B1125 |
|  | Phaseolus Vulgaris  Leucoagglutinin | B1115 |

Fig. S1. Phenotypic Characterisation of mVSCs and C3H 10T1/2 cells

**A.** Representative immunocytochemical images of mVSCs and C3H310T1/2 cells stained with DAPI, anti-S100β, anti-Sca1, anti-Cnn1 or anti-Myh11. MOVAS SMCs were used as a positive control for SMC differentiation markers (Cnn1 and Myh11). Control cells stained with secondary antibody alone was used to determine level of background/off-target fluorescence. Scale bar = 25μm. Data are representative of 10 images from n=3.

**B.** The relative telomere length of genomic DNA isolated from fresh aortic smooth muscle cells, SMCs cultured *in vitro* and S100β mVSCs isolated from the aortic arch using real time PCR. Amplification was relative to a single copy reference (SCR) gene. Data are presented as the ratio of telomere length relative to the aortic tissue SMC sample. Data are the mean ± SEM and representative of n=3 *p ≤ 0.05

**C.** The level of telomerase activity in protein lysates from aortic tissue and S100β mVSCs. Amount of PCR product formed in each sample was compared against a standard curve to determine level of telomerase activity which is given as total protein generated (TPG) units. Data are the mean ± SEM and representative of n=3 *p ≤ 0.05

**D, E.** Real-time qRT-PCR analysis of the relative expression of adipogenic marker genes (*Lpl* and *Fabp4*) and osteogenic markers (*Alpl* and *Sost*) in S100β mVSCs treated for 14 days and 21 days with adipogenic and osteogenic differentiation media, respectively. The housekeeping gene, hypoxanthine phosphoribosyltransferase (*hprt)* was used as control. Data are the mean ± SEM and representative of n=3 *p ≤ 0.05

**Fig. S2. Transfection Efficiency of siRNA and optimisation of Notch1 immunoprecipitation**

**A.** Cytochemical microscopic analysis of S100β mVSCs transfected with Tye563 DsiRNA for 48 h. DAPI was used to stain nuclei. Tye563 positive cells were counted and compared against total cell count. Representative fluorescence images of Tye563-Red expression in both transfected and non-transfected samples. Scale bar = 50μm

**B, C.** Expression of Notch1 protein in S100β mVSCs following *Notch1* siRNA knockdown. Cells were transfected with Notch1 siRNA (25nM) and scrambled control for 48 h at 37^o^C in MM1 before Notch1 protein was assessed in cell lysates by immunoblot. (B) A representative immunoblot of Notch1 at 120kDa and β-actin at 42kDa. (C) Densitometric analysis of the intensity of Notch1 band was normalised to the β-actin loading control. Notch1 protein expression is presented as the ratio of expression relative to the negative control sample transfected with non-targeted (scrambled) siRNA. Data are the mean ± SEM and representative of n=3 *p ≤ 0.05

**D.** Real-time qRT-PCR analysis of the relative expression of *Notch1* mRNA levels in S100β mVSCs following *Notch1* siRNA knockdown relative to the scrambled control. Cells were transfected with Notch1 siRNA (25nM) and scrambled control for 48 h at 37^o^C in MM1. The housekeeping gene, hypoxanthine phosphoribosyltransferase (*hprt)* was used as control. Data are the mean ± SEM and representative of n=3 *p ≤ 0.05

**E.** Expression of Notch1 protein purified from S100β mVSCs lysates following immunoprecipitation with protein A-conjugated magnetic beads. A representative immunoblot image of Notch1 bands at 120kDa. All other bands represent off-target staining.

**Fig. S3.** **Ectopic expression of Notch1 and Notch1 NICD in S100β mVSCs and the effect of chemical inhibition of N-glycosylation and *Mgat3* and *Fut8* knockdown on Notch1.**

**A.** Ectopic expression of myc-tagged Notch1 and NICD in S100β mVSCs. Cells were transfected with either full length Notch1 plasmid, Notch1 NICD plasmid or an empty vector and incubated for 48 h at 37^o^C in MM1 before expression of myc-tagged Notch1 and NICD was assessed by immunoblot. Representative immunoblot image shows full-length Notch1 bands at ~300kDa, NICD bands ~ 88kDa and β-actin bands at ~ 42kDa.

**B.** Densitometric analysis of the intensity of Notch1 and NICD bands normalised to the β-actin loading control. Full-length Notch1 and NICD protein levels as are expressed as a ratio of expression relative to the empty vector. Data are the mean ± SEM and representative of n=3 *p ≤ 0.05.

**C.** Real-time qRT-PCR analysis of the relative expression of *Notch1* mRNA levels in S100β mVSCs following treatment of cells with TNC (0.5μg/mL) or DMJ (4μg/mL) for 48 h at 37^o^C. Data are presented as the ratio of expression relative to the untreated negative control sample. The housekeeping gene, hypoxanthine phosphoribosyltransferase (*hprt*) was used as control. Data are the mean ± SEM and representative of n=3 *p ≤ 0.05.

**D-E.** Notch1 protein expression in S100β mVSCs following treatment of cells with TNC (0.5μg/mL) or DMJ (4μg/mL) for 48 h at 37^o^C. (D) A representative immunoblot image of Notch1 bands at 120kDa and β-actin bands at 42kDa. (E) Densitometric analysis of the intensity of Notch1 normalised to the β-actin loading control. Notch1 protein levels given as ratio of expression relative to the untreated negative control sample. Data are the mean ± SEM and representative of n=3 *p ≤ 0.05.

**F-G.** Real-time qRT-PCR analysis of the relative expression of (F) *Mgat3* and (G) *Fut8* mRNA levels in S100β mVSCs following *Mgat3* and *Fut8* knockdown. Data are presented as the ratio of expression relative to the scrambled control sample. The housekeeping gene, hypoxanthine phosphoribosyltransferase (*hprt*) was used as control. Data are the mean ± SEM and representative of n=3 *p ≤ 0.05.

Fig. S4. Chemical inhibition of N-glycosylation (TNC and DMJ) on lectin binding and Jagged-1 activation of Notch1 signalling and SMC differentiation in C3H10T1/2 cells.

**A-B.** Lectin binding analysis of N-glycans on whole cell lysates from C3H10T1/2 cells following treatment of cells with a range of (A) TNC and (B) DMJ concentrations for 48 h at 37^o^C. Data are presented as the mean absorbance value at 450 nm normalised to the negative control TBST sample. Data are the mean ± SEM and representative of n=3 *p ≤ 0.05 compared to the non-treated sample probed with the same lectins.

**C.** Lectin binding analysis of N-glycans on whole cell lysates from C3H10T1/2 cells following *Mgat3* and *Fut8* knockdown for 48 h at 37^o^C. Data are presented as the mean absorbance value at 450 nm normalised to the negative control TBST sample. Data are the mean ± SEM and representative of n=3 *p ≤ 0.05 compared to the scrambled control probed with the same lectins.

**D-E.** Real-time qRT-PCR analysis of *Hey1* (D) and *Myh11* (E) mRNA levels in C3H10T1/2 cells treated with recombinant immobilised Jagged-1 (Jag-1 Fc) or IgG-Fc (Fc) at 1μg/mL in MM1 media after 48 hrs following prior treatment of cells with TNC (0.5μg/mL). The housekeeping gene, hypoxanthine phosphoribosyltransferase *(hpr*t) was used as control. Data are the mean ± SEM and representative of n=3 *p ≤ 0.05

**F-G.** Real-time qRT-PCR analysis of *Hey1* (F) and *Myh11* (G) mRNA levels in C3H10T1/2 cells treated with recombinant immobilised Jagged-1 (Jag-1 Fc) or IgG-Fc (Fc) at 1μg/mL in MM1 media after 48 hrs following prior treatment of cells with DMJ (4μg/mL). The housekeeping gene, hypoxanthine phosphoribosyltransferase *(hpr*t) was used as control. Data are the mean ± SEM and representative of n=3 *p ≤ 0.05.

**Fig. S5.** ***Mgat3* and *Fut8* knockdown on Jagged-1 activation of Notch1 signalling and SMC differentiation in C3H10T1/2 cells.**

**A.** Cytochemical microscopic analysis of transfected C3H10T1/2 cells with Tye563 DsiRNA for 48 h. DAPI was used to stain nuclei. Tye563 red positive cells were counted and compared against total cell count. Representative fluorescence images of Tye563 expression in both transfected and non-transfected samples. Scale bar = 25μm.

**B.** Real-time qRT-PCR analysis of the relative expression of *Mgat3* and *Fut8* mRNA levels in C3H10T1/2 cells following *Mgat3* and *Fut8* knockdown. Data are presented as the ratio of expression relative to the scrambled control sample. The housekeeping gene, hypoxanthine phosphoribosyltransferase (*hprt)* was used as control. Data are the mean ± SEM and representative of n=3 *p ≤ 0.05.

**C-D** Real-time qRT-PCR analysis of *Hey1* (C) and *Myh11* (D) mRNA levels in C3H10T1/2 cells treated with recombinant immobilised Jagged-1 (Jag-1 Fc) or IgG-Fc (Fc) at 1μg/mL in MM1 media after 48 hrs following *Mgat3* knockdown. The housekeeping gene, hypoxanthine phosphoribosyl transferase (*hprt*) was used as control. Data are the mean ± SEM and representative of n=3 *p ≤ 0.05.

**E-F.** Real-time qRT-PCR analysis of *Hey1* (E) and *Myh11* (F) mRNA levels in C3H10T1/2 cells treated with recombinant immobilised Jagged-1 (Jag-1 Fc) or IgG-Fc (Fc) at 1μg/mL in MM1 media after 48 hrs following Fut8 knockdown. The housekeeping gene, hypoxanthine phosphoribosyl transferase (*hprt*) was used as control. Data are the mean ± SEM and representative of n=3 *p ≤ 0.05.

**Fig. S6. Puromycin selection and transfection efficiency in C3H10T1/2 cells**

**A.** Validation of puromycin selection in C3H10T1/2 cells. Cells were transfected with a range of concentrations of puromycin resistance-coding plasmid DNA ± puromycin (3μg/mL). Cells were fixed and nuclei stained with DAPI. Representative images of DAPI nuclei at 20X magnification. Scale bar = 25μm. Cells were counted using ImageJ and data graphed as mean cell number. Cell survival rates against puromycin were calculated for each concentration of plasmid DNA and compared to the equivalent non-puromycin-treated control. Data are the mean ± SEM and representative of n=3 *p ≤ 0.05.

**B.** Plasmid transfection efficiency in C3H10T1/2 cells. Cells were transfected with a range of concentrations of GFP-coding plasmid DNA and incubated for 48 h at 37oC. Cells were fixed and nuclei stained with DAPI and 10 images were analysed per sample. Representative images at 20X. Scale bar = 25μm. Cells counted using ImageJ and data graphed as percentage cell death compared to the non-transfected control. (c) GFP-positive cells counted and graphed as a percentage of total cell number. Data are the mean ± SEM and representative of n=3 *p ≤ 0.05 compared to non-transfected control.

**Fig. S7. Mutant plasmid sequence alignment and chromatograms validating the quality of sequencing reads at the site of each Notch1 N-glycosylation site mutation.**

**A.** Mutant plasmids were sequenced and aligned to wild-type Notch1 plasmid sequence using Multalin online sequence alignment tool. Image shows screenshots of the sections of the alignments containing the desired single base substitutions. Red letters indicate mutual consensus between wild-type and mutant plasmid. Blue or black letters indicate discrepancy between wild-type and mutant plasmids.

**B.** Chromatograms validating the quality of sequencing reads at the site of each Notch1 N-glycosylation site mutation.

**Fig. S8. Effect of asparagine mutations on plasmid transfection efficiency in** **C3H10T1/2 cells**

**A.** PCR analysis of the relative levels of Notch1-coding plasmid DNA in C3H10T1/2 cells co-transfected with puromycin resistance plasmid and either the wild-type Notch1 plasmid or one of the six mutant Notch1 plasmids and incubated for 48 h at 37oC, with puromycin treatment (3μg/mL) for the final 24 h. Data are presented as the ratio of expression relative to the wild-type control. Data are the mean ± SEM and representative of n=3 *p ≤ 0.05 compared to non-transfected control.

**B.** Plasmid Validation. NICD, Notch1, GFP and Puromycin (Puro) resistance plasmids were purified from bacterial cultures and digested with an appropriate restriction enzyme (BamHI or EcoRI) to linearise DNA. Digested and undigested samples were run on a 1% Agarose Gel and visualised under UV light to validate plasmid purity and length. Data representative of n=3. (Note: Incorrect restriction enzyme used for NICD plasmid resulting in incorrect plasmid length).

**C.** NICD Plasmid Validation. NICD Plasmid was purified from bacterial culture and digested with EcoRI restriction enzyme to linearise DNA. Digested and undigested samples were run on a 1% Agarose Gel and visualised under UV light to validate plasmid purity and length.
